# 5-Azacytidine incorporation into mRNAs disrupts translation and induces ribosome collisions

**DOI:** 10.64898/2026.03.30.714548

**Authors:** Alexis B Roberson, James Marks, Ruby Pitts, Bavavarshini Tamilselvam, Brian Grieb, William P Tansey, Sezen Meydan

**Author notes:** Greco-Hainsworth Centers for Research, Tennessee Oncology & OneOncology, Nashville, TN 37203. Contributed equally (Alexis B Roberson, James Marks).

## Abstract

5-Azacytidine (5-AzaC) is a cytidine analog and is widely used to treat myelodysplastic syndromes (MDS) and acute myeloid leukemia (AML). Although its therapeutic activity is primarily attributed to hypomethylation resulting from DNA incorporation, the majority of 5-AzaC is incorporated into RNA. However, the functional consequences of 5-AzaC incorporation into RNA have been unknown. Here, we show that 5-AzaC treatment of cells leads to inhibition of protein synthesis. Ribo-seq, Disome-seq, and RNA-seq in cells treated with 5-AzaC exhibit a time-dependent C-to-G transversion signature in mRNAs within 2 h of treatment. These transversion events are enriched within footprint positions corresponding to the A-site of monosomes or leading stalled ribosome in a disome complex. Consistently, ribosome and disome footprints are accumulated at sites with C-rich codons in the A-site, specifically with the codons containing a C in the second position. 5-AzaC activates the integrated stress response (ISR) and the ribotoxic stress response (RSR) in a GCN2- and ZAK-dependent manner, consistent with disome-mediated signaling. Furthermore, loss of the Ribosome Quality Control (RQC) factor, ZNF598, sensitizes cells to 5-AzaC. Collectively, our results support a model where 5-AzaC is rapidly incorporated into mRNAs, disrupts decoding, and triggers disome-mediated signaling pathways, which contribute to its cytotoxicity. These findings suggest that translation disruption represents an additional layer of 5-AzaC’s mechanism of action, alongside its known DNA-mediated effects.

## INTRODUCTION

Nucleic acids serve a vital function by storing and expressing genetic information. This functionality is housed within the common chemical structure of nucleotides composed of a five-carbon sugar, phosphate group, and nitrogenous base. However, the critical chemistry enabled by these structures is susceptible to damage by exogenous or endogenous insults such as oxidative stress, ultraviolet (UV) irradiation, and alkylating agents^1-3^, resulting in adduct formation, crosslinking, strand breaks, and the formation of abasic sites^4^. Although these insults can damage both DNA and RNA, the primary focus of the compromised nucleic acid integrity has been centered on DNA damage based on the assumption that DNA damage leads to permanent, long-term changes to genetic information while RNA damage can be mitigated through degradation of damaged RNA species^3,4^. However, this assumption is challenged by the long-lived nature of functional RNAs, including rRNAs^5^ and tRNAs^6^, as well as the relatively stable half lives of many eukaryotic mRNAs^7,8^. Furthermore, the single-stranded nature and primarily cytoplasmic localization of RNA further increase its vulnerability to damage compared to double-stranded DNA, which is protected in the nucleus by chromatin organization^9-11^. In line with this, RNA damage can disrupt cellular processes^12-17^ and has been linked to diseases^9,18-21^.

One way cells survey and respond to mRNA damage is mediated through ribosomes. Many forms of mRNA damage, such as oxidation, alkylation, or crosslinking of nucleotides, disrupt translation elongation, resulting in ribosome stalling at these sites, followed by collisions with trailing ribosomes^15-17,22-26^. The collided ribosome complexes, also referred to as disomes, form a unique interface that serves as a platform for recognition by quality control and stress response pathways^27-30^. The ribosome-associated quality control (RQC) is mediated by the E3 ubiquitin ligase ZNF598 (Hel2 in yeast), which promotes the ubiquitination of small ribosomal subunit proteins to promote ribosome rescue and nascent peptide degradation^31-34^. When disome levels exceed the capacity of the RQC to resolve them, disomes activate the Integrated Stress Response (ISR) and Ribotoxic Stress Response (RSR) pathways that are aimed at promoting cell survival or triggering cell death^12,16,17,35,36^. Ribosome collisions trigger activation of the ISR through phosphorylation of the eukaryotic initiation factor eIF2α via the kinase GCN2 and its co-activator GCN1^17,35-37^, resulting in global inhibition of cap-dependent translation alongside selective translation of stress-responsive transcripts such as *ATF4*^38,39^. RSR activation occurs through the recruitment of the MAP3 kinase ZAK to collided ribosomes, which activates the stress-activated protein kinases, p38 and JNK, leading to cell cycle arrest and apoptosis^12,16,17,40,41^. Together, these pathways position ribosome collisions as a signaling hub that senses translation stress and determines cellular fate. However, how different sources of RNA damage engage these pathways and ultimately influence cell fate is not fully understood.

Despite the growing understanding of how RNA damage is sensed at the level of translation, its role in clinically relevant contexts remains underexplored. A key gap lies in the context of chemotherapeutic agents that are classified as nucleobase and nucleoside analogs. Like their counterparts derived from nucleic acid damage, these analogs have altered physiochemical structures as compared to the endogenous nucleotides. These compounds are traditionally thought to exert their effects through incorporation into DNA and disruption of DNA metabolism^42^. However, many of these analogs also incorporate into RNA and perturb RNA-dependent processes, raising the possibility that RNA damage contributes to their cytotoxicity^42^. A prominent example is 5-fluorouracil (5-FU), a widely used pyrimidine analog for the treatment of colorectal, pancreatic, and breast cancers^43^. 5-FU was originally proposed to act primarily through the inhibition of thymidylate synthase, thereby impairing DNA synthesis^43,44^. However, several studies demonstrated that 5-FU-induced RNA damage disrupts several RNA-mediated processes^45-51^. Notably, 5-FU incorporation into RNA was shown to induce ribosome collisions and activate the RQC pathway^52^. These findings challenge the prevailing DNA-centric view of chemotherapeutic mechanisms and highlight RNA damage as an important determinant of drug response.

Another example of a nucleotide analog that is similarly classified as a DNA-targeting agent is a cytidine analog, 5-Azacytidine (5-AzaC), which is used in the treatment of acute myeloid leukemia (AML) and myelodysplastic syndromes (MDS)^53^. Upon cellular uptake, the majority of 5-AzaC (approximately 80–90%) is incorporated into RNA, while only a minor fraction is converted by ribonucleotide reductase into a deoxy-form for DNA incorporation^54^. Nevertheless, the therapeutic mechanism of 5-AzaC has been extensively characterized in terms of DNA hypomethylation, driven by covalent trapping of DNA methyltransferases and subsequent genome-wide demethylation that would ultimately enhance the expression of tumor suppressor genes^53^. On the other hand, 5-AzaC was shown to impair methylation of mRNAs^55^, to reduce tRNA^Asp^ methylation^56^ and to inhibit non-sense mediated decay (NMD)^57^, hinting its link to RNA damage. However, it is unknown how 5-AzaC’s RNA incorporation disrupts downstream pathways and how RNA damage contributes to its cytotoxicity.

Here, we investigate the consequences of 5-AzaC treatment on protein synthesis and cellular stress. We find that 5-AzaC is preferentially incorporated into mRNAs with short half-lives and disrupts protein synthesis. Treatment with 5-AzaC results in ribosome stalling and collisions, and these stalled ribosomes are enriched when C-containing codons are at the A-site. Furthermore, 5-AzaC leads to the activation of disome-mediated surveillance pathways including RQC, ISR, and RSR mediated by ZNF598, GCN2, and ZAK, respectively. Together, these results along with other recent findings^58^ provide evidence that RNA damage and translational stress contribute to the cytotoxic mechanisms of 5-AzaC. More broadly, our results expand understanding of how nucleoside analogs perturb mRNAs and highlight disome signaling as a key mediator of chemotherapeutic stress.

## RESULTS

### 5-AzaC impairs translation and activates stress signaling

5-AzaC is a cytidine analog that contains a nitrogen at position 5 of the pyrimidine ring (Figure 1A). In cells, it is metabolized to 5-Aza-CTP and predominantly incorporated into RNA during transcription (Figure 1B), whereas a smaller fraction is converted via ribonucleotide reductase to 5-Aza-dCDP to be incorporated into DNA (Figure 1B)^54^. Given its preferential incorporation into RNA, we hypothesized that 5-AzaC would disrupt protein synthesis by interfering with translation. To test this, we treated HCT116 cells with 5-AzaC and monitored global protein synthesis using puromycin incorporation assays. Treatment of cells with increasing concentrations of 5-AzaC for 8 hours (h) resulted in a dose-dependent inhibition of protein synthesis, with 10 µM 5-AzaC causing a substantial suppression and 100 µM leading to near-complete inhibition (Figure 1C). Based on these results, we selected 10 µM 5-AzaC to examine the kinetics of translation inhibition. Time-course experiments revealed that 5-AzaC treatment rapidly reduces protein synthesis (Figure 1D). A decrease in puromycin incorporation was already evident within 2h of treatment, and translation was nearly completely inhibited by 4-8h. Together, these findings demonstrate that 5-AzaC rapidly impairs protein synthesis in mammalian cells, consistent with the idea that its incorporation into mRNAs interferes with translation.

**Figure 1.**
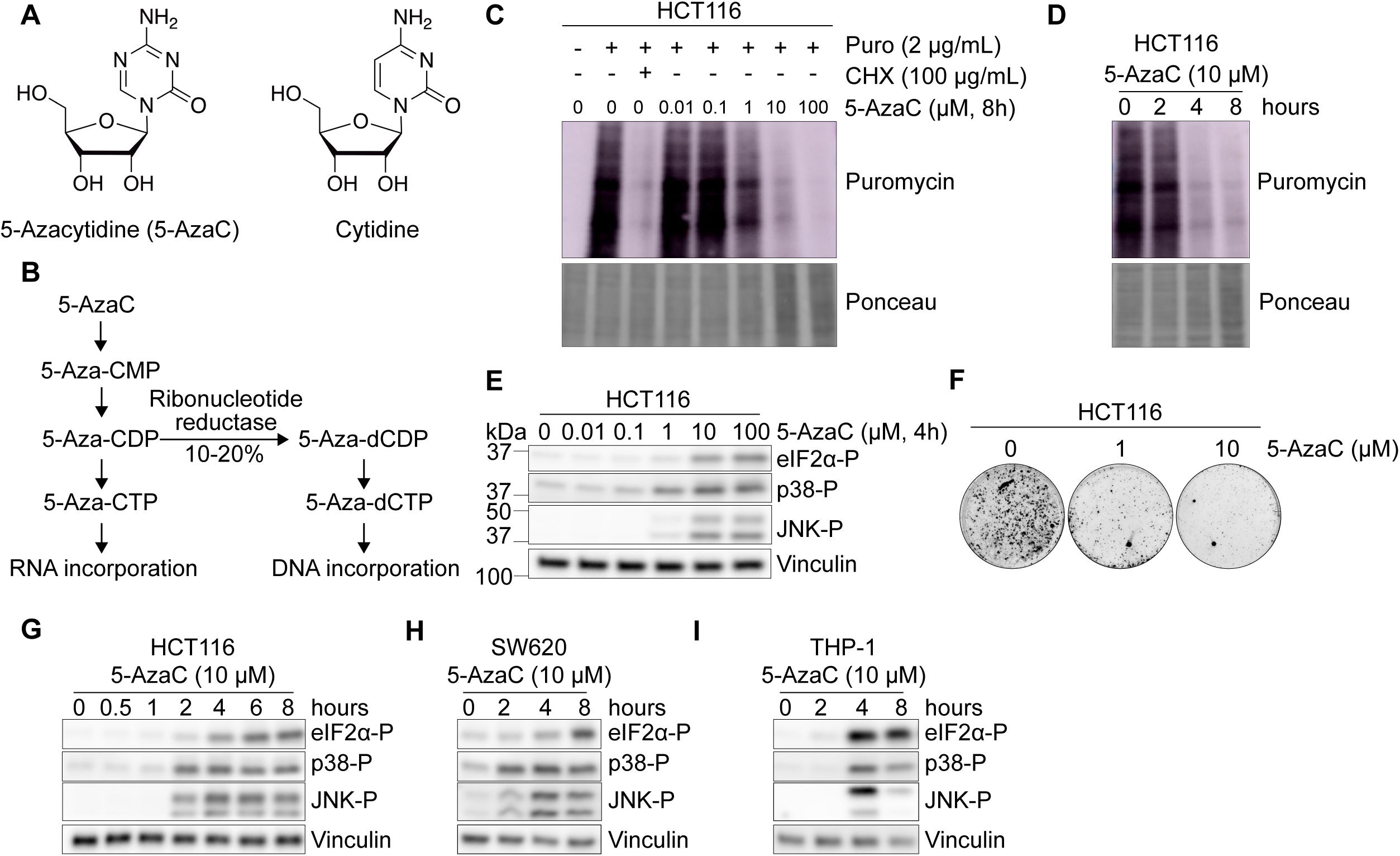
5-AzaC disrupts translation and activates stress response signaling. (A) Chemical structures of 5-Azacytidine (5-AzaC) versus cytidine, highlighting substitution of nitrogen for carbon at position 5 of the pyrimidine ring. (B) Schematic of 5-AzaC metabolism in the cell. Approximately 10-20% of 5-Aza-CDP is converted to 5-Aza-dCDP and incorporated into DNA, whereas 80-90% is incorporated into RNA. (C) Puromycin incorporation assay measuring protein synthesis in HCT116 cells treated with the indicated concentrations of 5-AzaC for 8 h. Cycloheximide (CHX) serves as a translation inhibition control. Ponceau staining is loading control. (D) Time course of puromycin incorporation assay in HCT116 cells treated with 10 µM 5-AzaC for 0-8 h. Ponceau staining is loading control. (E) Representative immunoblots of eIF2α phosphorylation (eIF2α-P), p38 phosphorylation (p38-P), JNK phosphorylation (JNK-P) in HCT116 cells treated with indicated concentrations of 5-AzaC for 4 h. Vinculin serves as loading control. (F) Colony formation assay in HCT116 cells with indicated concentrations of 5-AzaC. (G) Time course immunoblots of eIF2α-P, p38-P, JNK-P in HCT116 cells treated with 10 µM 5-AzaC for 0-8 h. Vinculin serves as loading control. (H) Time course immunoblots of eIF2α-P, p38-P, JNK-P in SW620 cells treated with 10 µM 5-AzaC for 0-8 h. Vinculin serves as loading control. (I) Time course immunoblots of eIF2α-P, p38-P, JNK-P in THP-1 cells treated with 10 µM 5-AzaC for 0-8 h. Vinculin serves as loading control. See also Figure S1.

Since the ribosome-mediated sensing of mRNA damage has been linked to stress signaling pathways, including the ISR and RSR^12,16,17^, we next tested whether 5-AzaC triggers their activation. To this end, HCT116 cells were treated with increasing concentrations of 5-AzaC, and pathway activation was assessed by immunoblotting for established signaling markers, phosphorylation of eIF2α (eIF2α-P, ISR), phosphorylation of p38 and JNK (p38-P and JNK-P, RSR). Robust activation of both ISR and RSR pathways was observed beginning at 1 µM 5-AzaC and increased further at 10 and 100 µM (Figures 1E, and S1A-B). Treatment with cytidine alone did not induce either pathway (Figure S1C), indicating that the stress signaling is specific to 5-AzaC. At the same concentration that inhibited translation (Figure 1C) and induced stress signaling (Figure 1E), 10 µM 5-AzaC led to near-complete loss of cell viability, as measured by both colony formation (Figure 1F and S1D) and shorter-term viability assay (Figure S1E). Both ISR and RSR signaling were robustly induced as early as 2h post-treatment and reached maximal levels at approximately 4h (Figure 1G). Notably, this timing closely parallels the onset of translation inhibition (Figure 1D), suggesting a link between 5-AzaC-induced translational perturbation and stress pathway activation. These observations were also reproducible in other human cell lines, such as SW620 colorectal cancer cells (Figure 1H) and THP-1 leukemia cells (Figure 1I). Together, these results show that 5-AzaC rapidly activates stress signaling pathways in parallel with inhibition of protein synthesis, consistent with a model where mRNA incorporation of 5-AzaC disrupts translation and triggers ribosome-mediated stress responses.

### Inhibition of disome-sensing proteins disrupt 5-AzaC-induced stress signaling

We next sought to determine whether the ISR and RSR signaling observed upon 5-AzaC treatment is triggered by disome sensing. These pathways can be pharmacologically interrupted using the small molecule inhibitors A-92, which inhibits the ISR kinase GCN2, and Nilotinib, which inhibits the ZAK that mediates RSR in response to ribosome collisions^16^. Both inhibitors attenuated the stress signaling in HCT116 cells in the presence of disome-inducing concentrations of the translation inhibitor anisomycin^17^ (Figure S2A), confirming their activity in our system.

We first tested the role of GCN2 in mediating ISR activation following 5-AzaC treatment. Pre-treatment of cells with A-92 markedly reduced 5-AzaC-induced eIF2α-P, demonstrating that ISR activation is GCN2-dependent (Figures 2A and S2B). A-92 treatment also partially decreased phosphorylation of p38 and JNK, suggesting that ISR signaling contributes to RSR activation under these conditions. We also examined the role of ZAK in RSR activation. Treatment with Nilotinib abrogated 5-AzaC-induced phosphorylation of both p38 and JNK, indicating that RSR activation is ZAK-dependent (Figures 2A and S2B). Inhibition of ZAK by Nilotinib also reduced 5-AzaC-induced eIF2α-P, consistent with the finding that ZAK facilitates GCN2’s activity^17^. These responses were reproducible in THP-1 leukemia cells (Figure S2C), indicating that these signaling mechanisms are conserved across cell lines. Consistent with the pharmacological inhibition, knockout (KO) of ZAK in SW620 cells resulted in loss of 5-AzaC-induced phosphorylation of p38 and JNK (Figures 2B and S2D). We next aimed to determine how inhibition of disome sensing influences cell death in response to 5-AzaC. Interestingly, Nilotinib conferred resistance to 5-AzaC across all three cell lines tested, with a more pronounced effect in THP-1 cells (Figure S2E). However, this phenotype was not specific to ZAK inhibition, as increased resistance to 5-AzaC was also observed in SW620 ZAK KO cells (Figure S2F), suggesting an off-target effect. Consistent with its broad kinase inhibitory activity as a clinically used leukemia drug, Nilotinib may impact additional signaling pathways that modulate cellular sensitivity to 5-AzaC. Treatment with A-92 mildly reduced sensitivity to 5-AzaC (Figure S2E), in line with recent findings supporting a cytotoxic role for ISR activation in this context^58^.

**Figure 2.**
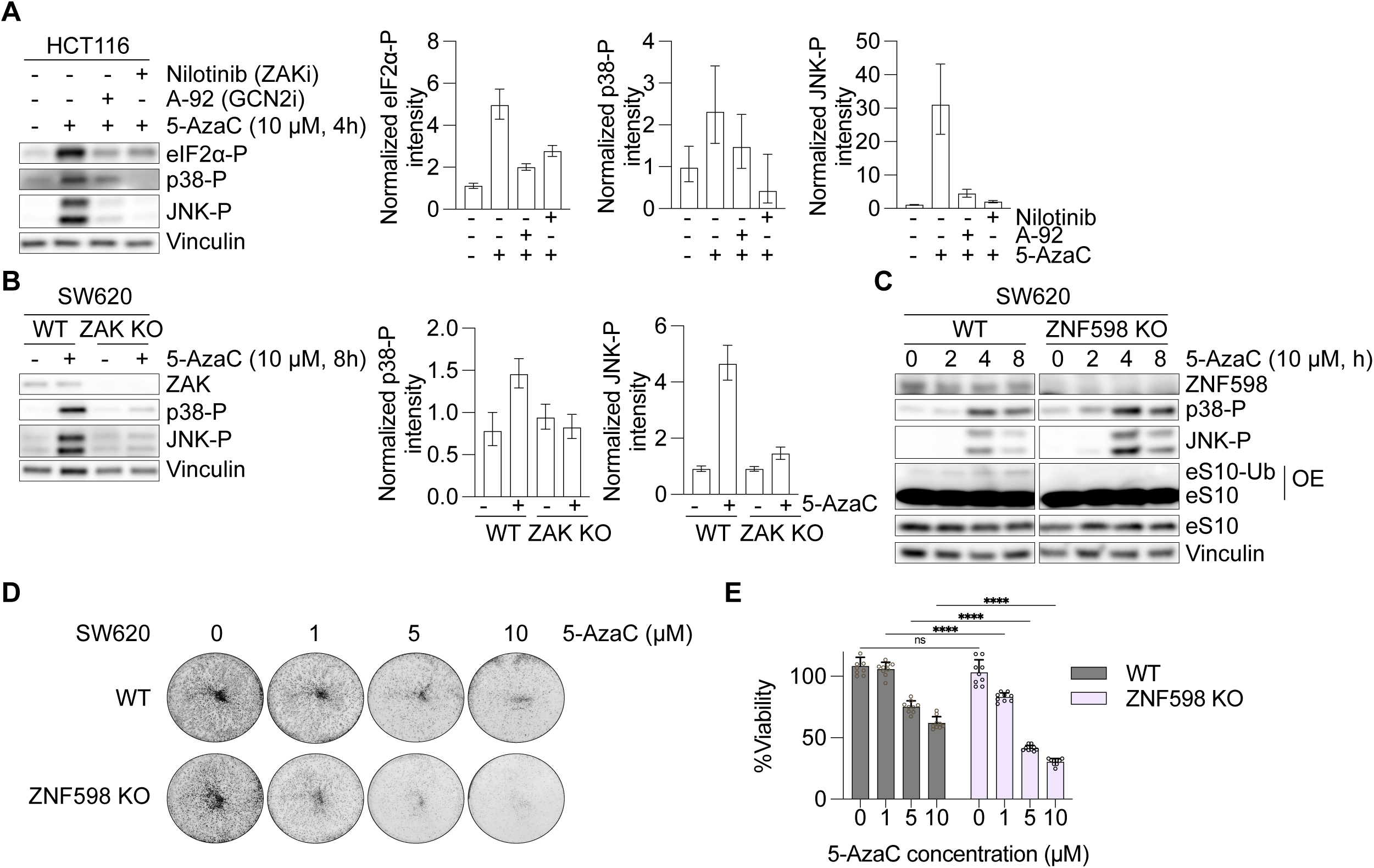
ISR and RSR signaling induced by 5-AzaC is dependent on GCN2 and ZAK. (A) Immunoblots of 5-AzaC-induced eIF2α-P, p38-P, JNK-P in HCT116 cells pre-treated with A-92 (2 µM, GCN2 inhibitor) or Nilotinib (1 µM, ZAK inhibitor) (left). Vinculin serves as loading control. Quantification of blots from three biological replicates is shown (right), n=3 biological replicates. (B) Immunoblots of ZAK, p38-P and JNK-P in SW620 WT and ZAK KO (left). Vinculin serves as loading control. Quantification of blots from three biological replicates is shown (right), n=3 biological replicates. (C) Immunoblots of ZNF598, p38-P, JNK-P, eS10 (ubiquitinated [eS10-Ub] and non-ubiquitinated forms) in SW620 WT and ZNF598 KO. Vinculin and eS10 serve as loading controls. Note that eS10 blot was overexposed to visualize eS10-ub. WT and ZNF598 KO samples were exposed together in the same blot, but later cropped for this image. (D) Representative colony formation assay (7 days) in SW620 WT and ZNF598 KO cells with indicated concentrations of 5-AzaC. (E) Viability assay (48 hours) measuring the effect of ZNF598 on 5-AzaC-induced cell. n=9 biological replicates. Statistical analysis was conducted with two-way ANOVA multiple comparisons test, ****=p<0.0001, ns=not significant (p=0.68). Where applicable, error bar indicates mean ± standard deviation. See also Figure S2.

Since we observed that 5-AzaC triggered ISR and RSR signaling in a GCN2- and ZAK-dependent manner, we next assessed whether RQC pathway is also engaged in response to 5-AzaC and how it contributes to its cytotoxicity. To this end, we monitored ribosomal protein eS10 ubiquitination, which is the target of ZNF598 upon disome recognition^59^. 5-AzaC treatment led to mild ubiquitination of eS10 beginning at 2h post-treatment, consistent with activation of disome-dependent RQC pathways (Figure 2C, eS10-Ub). eS10 ubiquitination was abolished in ZNF598 KO cells, confirming it is triggered by the RQC pathway (Figure 2C). Notably, loss of ZNF598 also enhanced p38 and JNK phosphorylation, suggesting that impaired RQC exacerbates disome-mediated stress signaling (Figure 2C). Consistent with this, we found that ZNF598 KO rendered cells more sensitive to 5-AzaC (Figures 2D-E and S2G). Taken together, these findings suggest that 5-AzaC incorporation into mRNA leads to ribosome collisions that activate canonical disome sensing pathways mediated by ZNF598, GCN2, and ZAK.

### 5-AzaC incorporation into mRNAs results in C-to-G conversions

To investigate the mechanisms by which 5-AzaC disrupts translation, and to detect 5-AzaC-induced gene expression changes, we conducted Ribo-seq, Disome-seq, and RNA-seq in HCT116 cells treated with 10 µM 5-AzaC for either 2 or 4h (Figure 3A). These time points were chosen based on our earlier observations that translation inhibition and stress signaling activation occur within this treatment window (Figures 1D and 1G). The RNA-seq analysis of differentially expressed genes revealed mild changes at 2h treatment but substantial changes with 4h of 5-AzaC (Figure 3B). Among these, significant upregulation of ISR and RSR transcriptional targets, such as *PPP1R15A*, *DDIT3*, *ATF3*, *JUN*, and *FOS,* were apparent (Figures 3B-C). Consistent with activation of the ISR, Ribo-seq showed a time-dependent increase in *ATF4* reads (Figure 3D), which is a canonical ISR target that is upregulated at the translation level^38,39^. These results indicate that 5-AzaC-induced stress signaling is consistently reflected in gene expression changes, which is more pronounced at 4h post-treatment.

**Figure 3.**
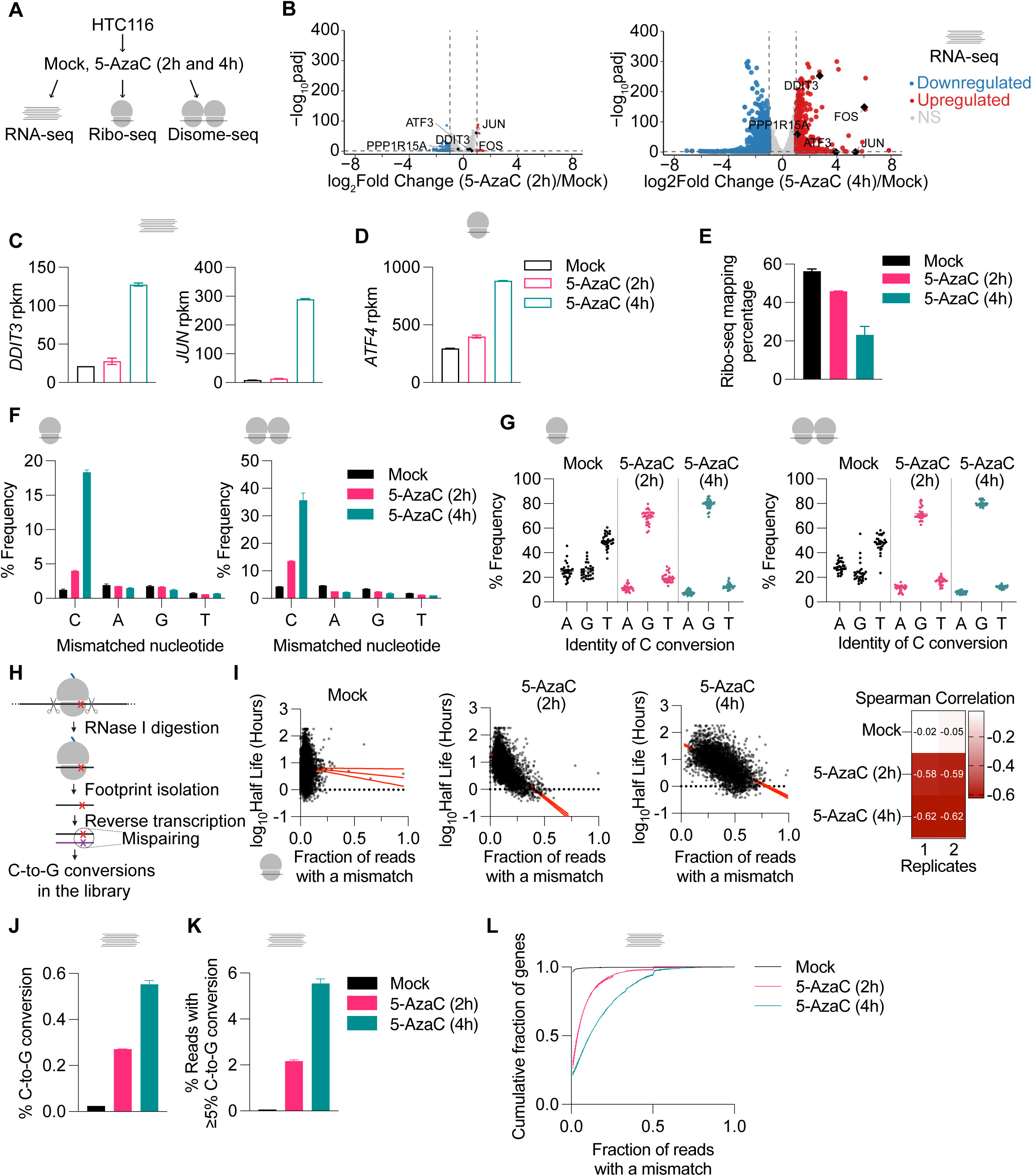
Incorporation of 5-AzaC into mRNAs causes C-to-G conversions. (A) Experimental design for RNA-seq, Ribo-seq and Disome-seq in HCT116 cells treated with 10 µM 5-AzaC for either 2h or 4h. Mock (DMSO)-treated cells act as controls. (B) Volcano plots of RNA-seq differential expression at 2h (left) or 4h (right) 5-AzaC treatment. Genes that are significantly upregulated (log2Fold Change > 1, adjusted p value [padj] < 0.05) or downregulated (log2Fold Change < -1, padj < 0.05) as determined by DESeq2 analysis are shown in red and blue, respectively. Genes with no significant changes (NS) are shown in gray. Individual genes are highlighted in diamond shape and are labeled. (C) RNA-seq expression levels of *DDIT3* and *JUN* over the course of 5-AzaC treatment. rpkm=reads per kilobase per million. (D) Ribo-seq expression levels of *ATF4* translation over the course of 5-AzaC treatment. rpkm=reads per kilobase per million. (E) Percentage of Ribo-seq reads mapping to the transcriptome over the course of 5-AzaC treatment. (F) Nucleotide mismatch frequencies for C, A, G, T in Ribo-seq (left) and Disome-seq (right) over the course of 5-AzaC treatment. (G) Identity of C conversion in Ribo-seq (left) and Disome-seq (right) footprints. Each point represents a nucleotide in the footprint (1-30 for Ribo-seq, 30-60 for Disome-seq). (H) Schematic of our library preparation process. Dashed circle is highlighting mispairing events occurring during reverse transcription that results in C-to-G conversions in the library. (I) Correlation between mRNA half-life and Ribo-seq mismatch rate in Mock (left), 5-AzaC (2h) (middle), and 5-AzaC (4h) (right). Representative data from a single replicate is shown, the other replicate can be found in Figure S3. Spearman correlation values are shown. (J) Percentage of C-to-G conversions in RNA-seq data. (K) Percentage of reads containing ≥5% C-to-G conversion in RNA-seq data. (L) Cumulative distribution of mismatch rates per gene in Mock and 5-AzaC treatment samples. Where applicable, error bar indicates mean ± standard deviation. See also Figure S3.

We next asked whether 5-AzaC’s incorporation into RNA leaves a detectable signature in our sequencing data. Indeed, during the initial processing of the Ribo-seq and Disome-seq datasets, we noticed that the total number of reads mapping to the transcriptome were lower in 5-AzaC-treated samples, especially at 4h post-treatment (Figures 3E and S3A). This reduction suggested that 5-AzaC treatment may introduce sequence mismatches that lower mapping efficiency. To test this, we quantified mismatch rates of each nucleotide across all mapped reads. In Mock-treated samples, only 1.2% of reads (average from 2 biological replicates) contained C mismatches in both Ribo-seq and Disome-seq datasets (Figure 3F). Strikingly, the fraction of C mismatches increased to 3.9% following 2h 5-AzaC treatment and further increased to 18.3% after 4h (Figure 3F). Notably, the mismatches observed in 5-AzaC-treated samples were almost exclusively associated with cytidines, with no detectable increase in mismatch rate at any other nucleotides (Figure 3F). Reads mapping to contaminating rRNA and tRNA species showed no elevation in mismatch rates, suggesting that C mismatches occur due to 5-AzaC’s incorporation into mRNAs (Figure S3B). To exclude the possibility that these mismatches arose from sequencing or library preparation artifacts, we added spike-in oligonucleotides (30mers and 60mers)^60^ matching the size of our monosome and disome footprints prior to the size selection step (Figure S3C). These spike-in oligos did not exhibit increased mismatch frequencies in 5-AzaC-treated sample library (Figure S3D). Although we did not specifically enrich for mitochondrial ribosome (mitoribosome) footprints, we still obtained reads mapping to mitochondrial mRNAs in our Ribo-seq data (Table S2). Our analysis of these reads revealed that 5-AzaC-induced C mismatches also occur in mitochondrial mRNAs (Figure S3E), indicating that 5-AzaC incorporation likely occurs across both cytosolic and mitochondrial transcripts. We next examined the identity of mismatch nucleotides in our dataset. In Mock-treated cells, background C mismatches were distributed across all three possible nucleotide conversions (Figure S3F). Notably, in 5-AzaC-treated samples, ∼70-80% of C mismatches corresponded to C-to-G conversions at 2-4h, with a smaller fraction of ∼10% converting to T (Figures 3G and S3G-H). This finding is consistent with previous data, where 5-AzaC was found to be irreversibly converted to ribosylguanylurea (RGU) via its pyrimidine ring opening in solution^61^. This open ring conformation was shown to inadvertently base pair with cytosine during reverse transcription first-strand cDNA synthesis, which leads to C-to-G conversions in the sequencing library^58,62^ (Figure 3H).

Having established that we can use mismatches as a tool to detect 5-AzaC’s incorporation, we next sought to determine which transcripts are prone to incorporate it. Because 5-AzaC is incorporated during transcription, we hypothesized that genes with higher transcription rate, as approximated by half-life, would have greater 5-AzaC incorporation. To test this, we calculated mismatch frequency per gene and integrated these values with a previously published RNA-seq dataset measuring mRNA half-lives in HCT116 cells using Actinomycin D transcriptional shutoff^63^. In Mock-treated samples, we observed no relationship between mismatch frequency and mRNA half-life (Figures 3I and S3I). On the other hand, after 2h of 5-AzaC treatment, a negative correlation emerged between mismatch frequency and transcript half-life (Figures 3I and S3I). By 4h of 5-AzaC, this anti-correlation became substantially stronger, where short-lived transcripts accumulated the highest mismatch rates, whereas long-lived transcripts remained relatively less affected (Figures 3I and S3I).

Finally, we aimed to quantify the extent of 5-AzaC incorporation into cellular RNAs. Since Ribo-seq and Disome-seq reads are short (<80 nt), allow only a single mismatch during mapping, and may be biased with footprints arising from ribosome stalling, we instead turned to our RNA-seq data for this analysis. RNA-seq provides longer reads (150 nt) and permits a higher mismatch threshold during mapping (up to 10 mismatches), enabling more sensitive detection of multiple incorporation events within individual transcripts. Consistent with our Ribo-seq and Disome-seq results, RNA-seq analysis revealed a time-dependent increase in C-to-G conversion rates following 5-AzaC treatment (Figure 3J). However, the average frequency of these events remained low: at 4h, only ∼0.54% of reads contained the C-to-G conversion compared to ∼0.024% in the Mock-treated samples. To estimate the fraction of mRNAs harboring 5-AzaC, we classified reads based on their individual rate of C-to-G mismatch and then applied a 5% cutoff. Only ∼5.5% of reads in the 4-hour 5-AzaC-treated sample met this criterion (Figure 3K), indicating that most mRNAs do not have a detectable 5-AzaC incorporation. We next extended this analysis to the gene level by applying the same cutoff to calculate per-gene incorporation rates and examining their cumulative distribution. At 2 and 4h, 50% of genes exhibited incorporation rates below ∼4% and ∼12%, respectively (Figure 3L). Taken together, these findings suggest that 5-AzaC is incorporated into newly synthesized mRNAs during transcription, preferentially affecting a subset of transcripts with shorter half-lives.

### 5-AzaC incorporation at A-site leads to ribosome stalling and collisions

Since 5-AzaC induces disome-mediated signaling (Figures 1 and 2) but is incorporated into a small subset of mRNAs (Figure 3), we next asked whether this incorporation is linked to ribosome stalling and collisions. To address this, we analyzed the Ribo-seq and Disome-seq footprints relative to the C mismatch positions. In Ribo-seq data, we observed a strong enrichment of ribosomes 16 nt downstream of the 5’ end of the footprint, corresponding to the second nucleotide of the A-site codon of the stalled ribosome in our dataset (Figure 4A). This enrichment was both time-dependent (Figure 4A), and specific to cytidines (Figure S4A). Consistently, Disome-seq data revealed a time-dependent enrichment of disomes 46 nt downstream of the 5’ end of the disome footprint (Figures 4B and S4B). This distance corresponds to the A-site of the leading, stalled ribosome in a disome complex, stacked with another ribosome behind that accounts for the additional 30 nt. These data indicate that 5-AzaC incorporation at the A-site codon is associated with ribosome stalling and subsequent disome formation. Ribosome stalling with an open A-site is known to generate shorter ribosome footprints^64^. Although we did not specifically enrich for these footprints, we nevertheless detected a time-dependent accumulation of 50-60 nt disome footprints in 5-AzaC-treated samples that were absent in Mock-treated samples (Figure 4C). In addition to the disome footprints getting shorter with increased 5-AzaC treatment, we also observed emergence of a 56-nt disome footprint at 4h. These changes in footprint length distribution could be reflective of 5-AzaC-induced disomes with open A-site.

**Figure 4.**
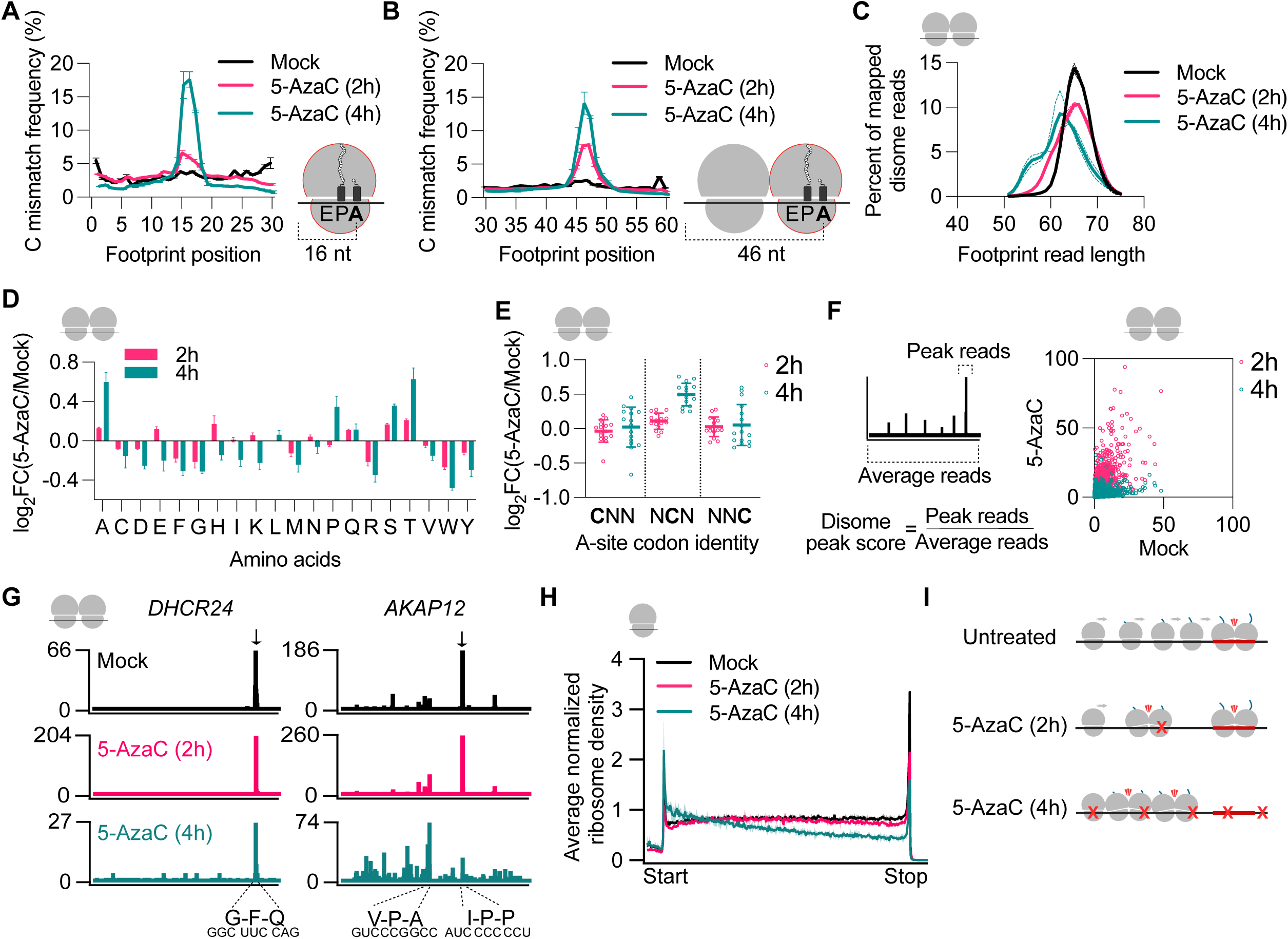
5-AzaC incorporation into mRNAs causes ribosome stalling and collisions. (A) C mismatch frequency across Ribo-seq footprint positions, showing 5-AzaC-induced enrichment at position 16, which corresponds to the A-site codon of the stalled ribosome. (B) C mismatch frequency across Disome-seq footprint positions, showing 5-AzaC-induced enrichment at position 46, which corresponds to the A-site codon of the stalled, leading ribosome in the disome complex. (C) Disome-seq footprint length distribution, showing accumulation of shorter footprints upon 5-AzaC treatment. Error bar is shown as dashed line. (D) Disome-seq pause score analysis of all amino acids showing 5-AzaC-induced enrichment at Ala, Pro, Ser, Thr codons. FC=Fold Change. (E) Disome-seq pause score analysis of C-containing codons at the A-site, demonstrating position-dependent effects with strongest stalling at the middle nucleotide. FC=Fold Change. (F) Schematic of disome peak score calculation (left) and distribution of disome peak scores in 5-AzaC (2h) and 5-AzaC (4h) samples compared to Mock-treated data. Average of two biological replicates is shown. (G) Disome-seq snapshots of *DHCR24* and *AKAP12* genes. Baseline disomes present in Mock-treated data are indicated with black arrows (top row), occurring at G-F-Q (*DHCR24)* and I-P-P *(AKAP12)* sites. These baseline disome peaks remain with 2h 5-AzaC treatment and get relatively more enriched (middle row, note the scale change), but are diminished or lost in 4h 5-AzaC treatment accompanied by emergence of new peaks (bottom row, note the scale change). 5-AzaC-induced disome site is highlighted in *AKAP12* gene that codes for V-P-A (GCC codon at the A-site). (H) Ribo-seq metagene analysis showing minimal changes in ribosome distribution with 2h 5-AzaC treatment but major re-distribution of ribosomes with 4h treatment, where ribosomes are progressively depleted from the 3’ regions of genes. (I) Model of 5-AzaC-induced translation effects across treatment course. At 2h, some disomes that were present in Mock-treated cells remain on the mRNAs as 5-AzaC incorporation is scarce. At 4h, 5-AzaC is incorporated at a higher frequency, which causes disome formation causing depletion of ribosomes from the 3’ regions of the mRNAs. The disomes that were present in Mock-treated cells are also lost during this re-distribution.

We next investigated the sequence context of 5-AzaC-induced ribosome stalling. Analysis of average pause scores across all amino acids revealed enrichment at codons encoding Pro, Ala, Ser, and Thr in 5-AzaC-treated samples (Figures 4D and S4C-D). Interestingly, these codons all have a C at their second positions. To directly test the effect of C position, we analyzed ribosome stalling at the codon level and found that having a C in the middle of the codon strongly promotes ribosome stalling and collisions, whereas C at the first or third positions have minimal impact (Figures 4E and S4E). These results indicate that the second position of the A-site codon is a key determinant of the 5-AzaC-associated ribosome stalling.

To identify gene-specific ribosome stalling events, we applied a disome peak scoring algorithm that quantifies peak intensity as the ratio of reads at each peak to the average reads across the corresponding transcript (Figure 4F). Analysis of the disome peak scores revealed unique disome re-distribution profiles upon 5-AzaC. At 2h, disome peaks that were present in Mock-treated samples were persistent and even relatively more enriched, while they were diminished or fully lost after 4h of 5-AzaC treatment (Figure 4F). Inspection of individual transcripts revealed a similar pattern, with prominent disome peaks in Mock-treated samples persisting at 2h and then becoming attenuated or lost at 4h (Figure 4G). To assess these effects more globally, we performed metagene analysis of Ribo-seq and Disome-seq data. While 2h of treatment led to minor changes, 4h treatment resulted in a pronounced re-distribution of ribosomes towards the 5’ ends of transcripts, with a corresponding depletion toward the 3’ ends (Figures 4H and S4F). A similar metagene profile was also observed with anisomycin treatment in human cells^65^, which prompted to ask whether this could be a pleiotropic effect of 5-AzaC-induced translation inhibition or if it is due to its incorporation into mRNAs. To address this, we separated our Ribo-seq data into four quartiles based on the mismatch rates of the genes and conducted metagene analysis for each quartile. We observed that the progressive ribosome depletion profile occurs only with genes carrying highest fraction of mismatches (Figures S4G and S4H, fourth quartile). Overall, these results support a model where 2h treatment leads to disome formation at relatively infrequent 5-AzaC-incorporated codons while baseline disomes at other codons remain unchanged (Figure 4I). In contrast, more extensive 5-AzaC incorporation at 4h induces widespread A-site ribosome stalling, leading to re-distribution of disomes and progressive depletion of ribosomes from downstream regions of the transcripts (Figure 4I).

## DISCUSSION

Nucleotide analog drugs such as 5-AzaC have long been used in the clinic, yet their effects on RNA-related processes remain poorly understood. Here, we show that 5-AzaC is incorporated into mRNAs, perturbs translation elongation, leads to ribosome stalling and collisions, and activation of disome-mediated signaling pathways (Figure 5). A key finding of this study is the identification of a C-to-G conversion signature in our sequencing data, which enables detection of 5-AzaC incorporation at single-nucleotide resolution and provides a quantitative readout of its global frequency. Although our data suggests that 5-AzaC incorporation preferentially affects only a subset of transcripts with shorter half-lives, the resulting RNA damage is sufficient to induce disome-mediated signaling and globally inhibit translation (Figure 5). This result is consistent with previous findings, where the transfection of a disome-inducing poly-Lys reporter or UV-treated EGFP mRNA was enough to elicit disome-mediated signaling^17^.

**Figure 5.**
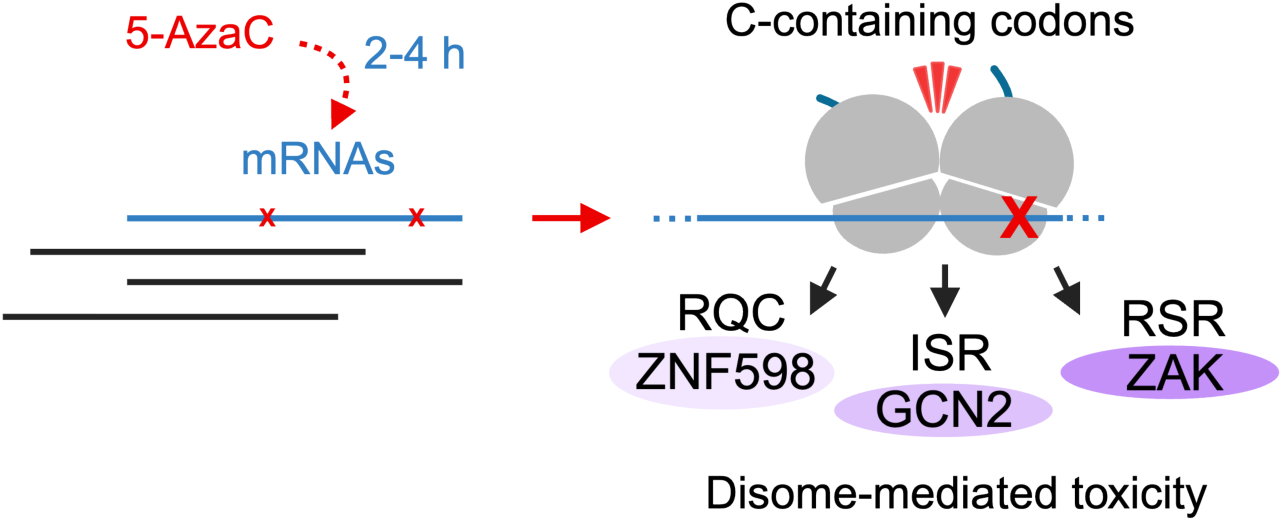
Model of 5-AzaC-induced ribosome stalling and collisions. We propose that 5-AzaC is incorporated into a subset of mRNAs (shown in blue) with shorter half-lives within the 2-4 hour treatment window of our experiments, whereas existing mRNAs (shown in black) are not affected. This incorporation is sufficient to induce ribosome stalling and collisions when the 5-AzaC-incorporated, C-containing codon is at the A-site of the ribosome. Disome-mediated signaling pathways, RQC, ISR, and RSR, are activated in response to 5-AzaC treatment and contribute to the 5-AzaC-induced cell death.

We found that 5-AzaC-induced C mismatches are strongly enriched at the A-site of the stalled ribosomes (Figures 4A-B). Furthermore, shorter disome footprints, which may be reflective of ribosomes with open A-site, accumulate during prolonged 5-AzaC treatment (Figure 4C). These findings suggest that 5-AzaC incorporation interferes with decoding, primarily affecting the C-containing codons for Ala, Pro, Ser, and Thr (Figure 4D). These codons are decoded by tRNAs containing inosine at position 34, modified by ADAT enzyme^66^. 5-AzaC could potentially impair ADAT activity by binding to its cytidine deaminase-like domain^67^. However, this is unlikely to explain our findings, as we did not observe exacerbated ribosome stalling at NNC/NCC codons (Figure S4E), which would be affected by lack of inosine modification, or at other codons decoded by inosine-containing tRNAs such as Leu, Ile, Val, and Arg (Figure 4D). Although 5-AzaC has been shown to inhibit methylation of tRNA^Asp^ after 24h of treatment^56^, we did not detect 5-AzaC-induced ribosome stalling at Asp codons (Figure 4E), likely due to our short treatment window, during which C-to-G conversions were not found in tRNAs and rRNAs (Figure S3B). Instead, C at the second position of the A-site codon emerged as a key determinant of 5-AzaC-induced ribosome stalling and collisions (Figure 4E). It remains unclear whether 5-AzaC itself directly interferes with decoding or the RGU formation specifically underlies this effect. Curiously, the same positional specificity was also observed with pseudouridine and *N*^1^-methylpseudouridine-incorporated mRNAs, where the middle nucleotide of the A-site codon exerts the strongest effect on ribosome stalling in the cell^68^, and on ribosomes reacting with near-cognate tRNAs in vitro^69^. Structural studies may elucidate the molecular basis by which 5-AzaC disrupts decoding at specific subcodon positions.

We propose that the mRNA damage is one of the mechanisms by which 5-AzaC induces cell death. Several findings from our study are consistent with this model, since we observed substantial disruption of translation (Figure 1C), induction of disome-mediated stress signaling (Figures 2A-C) and increased cell death in the absence of ZNF598 (Figures 2D-E). During the preparation of this manuscript, another study^58^ conducting a CRISPR/Cas9 screen with 5-AzaC found that loss of ZNF598 also resulted in increased sensitivity to the drug. One possible explanation for this finding could be that ZNF598 rescues ribosomes stalled at 5-AzaC-containing codons, and in its absence, ribosome collisions are exacerbated. Our data suggests that this could be the case, since we observed increased p38 and JNK phosphorylation in ZNF598 KO cells (Figure 2C). However, in contrast, loss of the downstream RQC factor ASCC3 that splits the stalled ribosomes, does not increase sensitivity of cells to 5-AzaC^58^, suggesting that lack of ribosome splitting may not fully explain the ZNF598 KO phenotype. It is possible that loss of ZNF598 impairs the rescue of baseline collisions, and thereby makes cells more sensitive to 5-AzaC. In addition to ZNF598, another ubiquitin ligase, RNF25, was found to play a critical role in 5-AzaC-induced cell death^58^. Interestingly, although UV-induced cell death was shown to occur via ZAK^16^, ZAK KO does not affect the sensitivity of cells to 5-AzaC^58^. Since ZAK-mediated p38 and JNK signaling was strongly activated in response to 5-AzaC-induced RNA damage (Figure 2B), the exact mechanisms by which cell death occurs and interplay between these different factors remain to be determined.

The clinical relevance of mRNA damage induced by 5-AzaC is an open question that needs to be investigated in future studies. Plasma concentrations of 5-AzaC following its administration to patients reach 1-5 µM^70,71^, which falls within the range where we observed the translation disruption in puromycin incorporation assays (Figure 1C). Given that 5-AzaC is typically administered over consecutive days, prolonged exposure may further increase its incorporation into mRNAs. It is unclear whether mRNA damage and disome formation occurs in the patients treated with 5-AzaC. The established mechanism of action of 5-AzaC is attributed primarily to DNA hypomethylation via DNA incorporation. In addition, deoxy version of the drug, 5-AzadC, which does not incorporate into RNA, is also an effective chemotherapeutic agent. Therefore, RNA damage alone is unlikely to account for the chemotherapeutic activity of 5-AzaC. However, RNA-mediated effects may contribute to its overall toxicity, particularly in non-dividing cells such as neurons^72,73^, where DNA incorporation could be limited. Interestingly, we observed incorporation of 5-AzaC to mitochondrial mRNAs (Figure S3E), raising the intriguing possibility that mitochondrial translation may also be perturbed. Whether this leads to mitoribosome stalling and contributes to mitochondrial dysfunction remains to be determined. Notably, inhibition of mitochondrial translation by tedizolid was shown to overcome the resistance to venetoclax^74^, a BCL-2 inhibitor drug that is used in a combination therapy with 5-AzaC for AML patients. If 5-AzaC impairs mitochondrial translation, co-inhibition with tedizolid could further exacerbate mitochondrial dysfunction and cellular stress, amplifying the activity of 5-AzaC-venetoclax combination therapy. These observations raise the possibility that mRNA damage-associated stress pathways, including those affecting mitochondrial translation, may influence both therapeutic response and toxicity, and could be leveraged or mitigated depending on clinical context.

Together, our findings establish that RNA damage-induced ribosome stalling and disome formation are underappreciated components of 5-AzaC activity. Ultimately, leveraging these insights may improve treatment strategies with existing drugs and can inform the design of next-generation nucleotide analogs that maximize anti-tumor efficacy while minimizing adverse effects.

## ACKNOWLEDGEMENTS

We are grateful to John Karijolich for the conception of the project, and his feedback on the manuscript. We also thank Katrin Karbstein for the critical reading of the manuscript, and Manny Ascano, Andrew Folkmann, and Karbstein Lab members for helpful discussions. We thank Rahul Bhowmick and David Cortez Lab for their help with colony formation assays. This study was supported by Fisk-Vanderbilt Bridge Program Fellowship to AR, P50CA236733 to WPT, and R00GM144688-04 to SM.

## AUTHOR CONTRIBUTIONS

SM and JM designed the study. ABR, JM, RP, BT, BG (under the supervision of WPT), and SM performed experiments. JM and SM conducted computational analysis. AR, JM, and SM wrote the manuscripts with input and edits from all authors.

## MATERIALS AND METHODS

### Cell lines

HCT116 cells (gift from Rahul Bhowmick) were cultured in McCoy’s 5A modified medium with *L*-glutamine (Gibco, 16600-082) supplemented with 10% fetal bovine serum (FBS) (Gibco, A56695-01). SW620 (gift from Robert Coffey) and THP-1 cells (gift from John Karijolich) were cultured in RPMI medium 1640 (Gibco, 11875-093) with *L*-glutamine supplemented with 10% FBS and an additional 2 mM *L*-glutamine (Gibco, 25030-081). HEK293T andHEK293FT cells (gift from Rahul Bhowmick) were cultured in DMEM (Gibco, 3199383) supplemented with 10% FBS. All cell cultures were grown at 37°C with 5% CO_2_ and tested for presence of mycoplasma contamination.

### Generation of SW620 “WT” cells expressing Cas9

To generate Cas9 expression lentivirus, HEK293T cells were transfected with the viral transfer plasmid plentiCas9-Blast (Addgene, 52962)^75^, the viral packaging plasmid psPAX2 (gift from Didier Trono; Addgene, 12260), and the viral envelope plasmid pMD2.G (gift from Didier Trono; Addgene, 12259) using Lipofectamine 3000 Transfection Reagent (Invitrogen, L3000015). After 48 hours, virus-containing media was collected and used with 8 µg/mL polybrene (Sigma, TR-1003) to transduce SW620 cells overnight. Following transduction, viral media was replaced with fresh media containing 10 µg/mL Blasticidin (Research Products International, B12200). A clonal SW620 Cas9 cell line was established by serial dilution of the population and screened for high Cas9 expression by Western blotting using mouse anti-Cas9 antibody (Cell Signaling 7A9-3A3, 14697S).

### Generation of SW620 ZNF598 and ZAK KO cells

For generating SW620 ZAK KO cells, ZAK-pLentiguide-Fwd (5’-CACCGTGTATGGTTATGGAACCGAG-3’) and ZAK-pLentiguide-Rev (5’-AAACCTCGGTTCCATAACCATACAC-3’) oligos were purchased from Integrated DNA Technologies. An oligo duplex was generated by incubating the oligos (100 μM each) with 10X T4 DNA ligase buffer (NEB, B0202) and T4 PNK (NEB, M0201) at 37°C for 30 minutes, followed by 95°C for 5 minutes, ramping down to 25°C at 5°C/min. The lentiGuide-Puro^75^ (Addgene plasmid # 52963) plasmid was digested using BsmB1-v2 restriction enzyme (NEB, R0739) and Buffer r3.1 (NEB, B6003), dephosphorylated by using CIP (NEB, M0525), and then gel purified. The digested vector was then ligated with 1:200 diluted oligo duplex by using T4 DNA ligase (NEB, M0202) by incubating at room temperature for 10 minutes. The ligation mixture was transformed to competent *E. coli* cells (MAX Efficiency STBL 2, Invitrogen, 10268-019). The resulting lentiGuide-Puro-ZAK plasmids were isolated from single colonies and their sequences were confirmed by Sanger sequencing with Genewiz via the U6 promoter. Lentivirus was produced as described above and was used to transduce SW620-Cas9 cells. Cells were maintained in 1.25 μg/mL puromycin (Selleck Chemicals, S7417) containing media for at least 1 week prior to knockout confirmation by western blotting and subsequent experiments.

For generating SW620 ZNF598 KO cells, ZNF598-pLentiguide-Fwd (5’-CACCGACCGCTGCTCTACCAAGATG-3’) and ZNF598-pLentiguide-Rev (5’-AAACCATCTTGGTAGAGCAGCGGTC-3’) oligos were purchased from Integrated DNA Technologies. An oligo duplex was generated by incubating the oligos (100 μM each) at 95°C for 5 minutes, then gradually ramping it down to 25°C in the presence of annealing buffer (100 mM potassium acetate, 30 mM HEPES-KOH pH 7.4, 2 mM magnesium acetate). The lentiGuide-Puro backbone was digested using BsmB1-v2 restriction enzyme (NEB, R0739) and Buffer r3.1 (NEB, B6003), and gel purified. 20 ng of the digested vector was ligated with 1:20 dilution of the oligo duplex via Ligation Mix (Takara, 6023) at 16°C for 30 minutes. The ligation mixture was transformed to NEB Stable Competent *E. coli* cells (NEB, C3040H). The resulting lentiGuide-Puro-ZNF598 plasmids were isolated from single colonies and their sequences were confirmed with Genewiz whole-plasmid sequencing. Lentivirus was produced in HEK293FT cells by transfecting them with lentiGuide-Puro-ZNF598 along with psPAX2, pMD2.G, pAdVantage (Promega, E1711) (gift from Michael Ward) as described above. 24h after transfection, the media was replaced with a fresh media containing 1:500 diluted Alstem ViralBoost Reagent (Fisher, NC0966705). After 48h, HEK293FT cells containing the lentiviral particles were collected by centrifugation at 1100xg for 5 minutes. Supernatant was filtered with a 0.22 μM filter and Alstem Lentivirus Precipitation Solution (Fisher, NC1080378) was added at a 1:3 dilution. This solution was kept at 4°C for 48h and then spun down at 4°C, 1500xg for 45 minutes. The supernatant was aspirated and the pellet was resuspended in 10x dilution of cold PBS (i.e. 600 μL of PBS per 6 mL media that was collected). 100 μL of this resuspension was added to SW620-Cas9 cells. Cells were maintained in 1.25 μg/mL puromycin containing media for at least 1 week prior to knockout confirmation by western blotting and subsequent experiments.

### Western blotting

Cells were grown to ∼80% confluency before treatment. All antibiotics, pharmacological inhibitors, and nucleoside analogues were prepared in DMSO. For concentration gradient and time course experiments, cells were treated with up to 100 μM 5-azacytidine (5-AzaC) (Millipore Sigma, A2385) for 4-12 hours or 10 μM 5-azaC for up to 8 hours prior to harvest, respectively. For GCN2 and ZAK inhibitor experiments, cells were pretreated with DMSO, 2 μM A-92 (Axon Medchem, 2720) or 1 μM Nilotinib (APExBio, A8232) for 30 minutes followed by treatment with 10 μM 5-AzaC for 4 hours or 1 μg/mL anisomycin (Selleck Chemicals, S7409) for 15 minutes before harvest. All subsequent experiments were performed with 10 μM 5-AzaC treatment unless otherwise stated.

For adherent cells, cell medium was aspirated and cells were washed once with ice-cold PBS. For suspension cells, the cell pellet was isolated by centrifugation at 500 x g for 5 minutes at 4°C and washed once with ice-cold PBS. All cells were lysed by incubation with RIPA lysis buffer (Thermo Fisher Scientific, 89900) supplemented with protease inhibitor (Sigma Aldrich, 4693159001) and phosphatase inhibitor (Selleck Chemicals, B15001 or Cell Signaling Technologies, 5870) on ice for 10 minutes and clarified via centrifugation at 12,000 rpm for 10 minutes at 4°C. Total protein concentration of clarified lysates was determined using the Pierce BCA protein assay kit (Thermo Fisher Scientific, 23225). Lysates were normalized to equal protein concentrations, and protein samples for blotting were prepared by mixing with Laemmli sample buffer (Bio-Rad, 1610737 or 1610747) supplemented with 2-mercaptoethanol (Sigma-Aldrich, M3148) and boiled for 5 minutes at 95°C. Proteins were resolved by SDS-PAGE on 4-20% Mini-PROTEAN TGX protein gels (Bio-Rad, 4561093 or 4561096) and transferred to PVDF membranes (Bio-Rad, 1704156) using the Trans-Blot Turbo transfer system. Membranes were blocked with 5% nonfat dry milk (Bio-Rad, 1706404) in TBST (blocking buffer) for 30 minutes at room temperature with gentle rocking. Primary and secondary antibodies were prepared as 1:1000 or 1:2500 dilutions in 5% blocking buffer, respectively. Blots were incubated with primary antibodies overnight at 4°C or for 2 hours at room temperature with gentle rocking. Blots were washed with TBST, 3 times for 5 minutes each, followed by incubation with secondary antibodies for 1 hour at room temperature with gentle rocking. Blots were then washed with TBST, 5 times for 5 minutes each, developed using Clarity Western ECL Substrate (Bio-Rad, 1705061) or SuperSignal West Femto Maximum Sensitivity Substrate (Thermo Fisher Scientific, 34094), and imaged on the Amersham Imager 600.

Antibodies used in this study: puromycin (Milipore Sigma, MABE343); eIF2α-P (Ser51) (Abcam, 32157); p38-P (Thr180/Tyr182) (Cell Signaling Technologies, 9211); JNK-P (Thr183/Tyr185) (Cell Signaling Technologies, 4668); ZAK (Bethyl Laboratories, A301-994A); ZNF598 (Abcam, 135921); RPS10 (Abcam, 151550); Vinculin (Abcam, 129002); H3 (Abcam, 10799); anti-rabbit IgG HRP Conjugate (Promega, W401B); anti-mouse IgG HRP Conjugate (Promega, W402B).

The band intensities of the blots were quantified by ImageJ2/Fiji version 2.16.0/1.54p. Background signal was first subtracted, then band intensities were measured using an identical selection area for all bands across the blot. The mean intensity of each band was normalized to the corresponding Vinculin loading control. The first sample was set to a value of 1, and all other values were normalized relative to this reference. The blots that were outputted from Fiji was cropped in Graphic Converter Version 12.2 to be placed in the figures.

### Puromycin incorporation assay

Protein synthesis was assessed using a puromycin incorporation assay. Puromycin and cycloheximide stocks were prepared in DMSO. Cells were treated with 2 μg/mL puromycin (Selleck Chemicals, S7417) 20 minutes prior to harvesting. For the cycloheximide control, cells were treated with 100 μg/mL cycloheximide (Sigma-Aldrich, C7698) 5 minutes prior to puromycin treatment. Lysates and protein samples for blotting were prepared as described above. After transfer of proteins to PVDF membrane, the membrane was washed for 2 minutes with Milli-Q H_2_O and stained with Ponceau S (Thermo Fisher Scientific, A40000279) for 10 minutes at room temperature with gentle rocking. Following staining, the membrane was washed 3-5 times with purified water for 3 minutes each, until red background was reduced and protein bands were clearly visible, and imaged on the Amersham Imager 600. The membrane was destained with 0.1% NaOH for 2 minutes, washed 3 times with purified water for 5 minutes each, and blocked with 5% blocking buffer for 30 minutes at room temperature with gentle rocking. Western blot analysis of nascent peptide synthesis using an anti-puromycin primary antibody (Millipore Sigma, MABE343) was performed as described above.

### Colony formation assay

Cells were seeded 400 per well on a 6-well plate with 2 mL of media. Immediately after seeding, the designated wells were treated with 5-AzaC treatment at indicated concentrations. 7 days after the seeding (once a single colony in Mock-treated well has >50 cells), the media was removed and each well was washed with PBS. 2 mL of 0.02% crystal violet dye (Sigma, C6158-50G) in 10% methanol was added to each well, and plates were rocked for 20 minutes. The dye was removed and each well was washed three times with purified water and left to dry before imaging with BioRad ChemiDoc system. The images corresponding to the well were cropped in Adobe Illustrator.

### Cell viability assay

Cells were seeded 2,000 per well on a 96-well plate in 100 µL of media. 5-AzaC, A-92, and Nilotinib were added at indicated concentration to each well 24h after seeding. 48h following treatment, CellTiter-Glo reagent (Promega #G7570) was added following manufacturer’s protocols and the samples were transferred to an opaque 96-well plate. Viability was measured by using BioTek plate reader to record luminescence.

### Ribo-seq and Disome-seq lysis, footprinting, and size selection

The cells are lysed as previously described^76^: A 10-cm dish of HCT116 cells were grown to ∼80% confluence and then treated with either DMSO for 4h or 10 µM 5-AzaC (Millipore Sigma, A2385) for either 2 or 4h. Cells were washed with chilled PBS and then flash frozen by floating the plates on liquid nitrogen. 400 µL of lysis buffer (1.3 mL polysome buffer (20 mM Tris-HCl pH 7.5, 150 mM NaCl, 5 mM MgCl_2_, 1 mM DTT, 100 µg/ml Cycloheximide(Sigma Aldrich, C1988-G), 150 µL 10% Triton X-100 (Sigma, T9284), 18.8 µL Turbo DNase (Fisher Scientific, AM2238)) was added to the frozen plates. Frozen cells in lysis buffer were thawed on wet ice, transferred to a 1.5 mL microfuge tube, and incubated on ice for 10 minutes. Lysate was triturated 10 times with a 26-G needle and RNA concentration was measured using a Qubit RNA BR Assay Kit (Fisher Scientific, Q10211). To recover both monosome and disome footprints^77^, lysate containing 40 µg of RNA was supplemented with lysis buffer to 300 µL and then digested with 80 U of RNase I (Thermofisher, AM2294) for 1 hour at 22°C shaking at 700 rpm. Digestion was stopped with 40 µl of SUPERase-In RNase inhibitor (Thermofisher, AM2696) and clarified by centrifugation at 21,000 x g at 4°C for 5 minutes. Samples were layered over 900 µL sucrose cushion buffer (4.68 mL polysome buffer, 2.04 g sucrose, 6 µL SUPERase-In) in 13 mm x 51 mm polycarbonate ultracentrifuge tubes. Samples were spun in a TLA100.3 rotor at 100,000 rpm for 1 hour at 4°C. Footprints were extracted by TRIzol (Fisher, 15596026)/chloroform and precipitated with isopropanol. Pellets were resuspended in 15 µL of 10 mM Tris-HCl, pH 8. Prior to size selection, 20 fmol of spike-in oligos were added to footprints. Size selection was carried out by resolving the samples on a 15% TBE Urea gel (Bio-rad, 3450091). RNA sizes markers for 25, 34, and 80 nt as well as an RNA ladder (Abnova, R0007) were run alongside samples. The gel was stained with SYBR gold for 5 minutes and monosome footprints (25-34 nt) and disome footprints (50-80) were excised from the gel along with RNA size markers and the 50 nt band from the RNA ladder. 700 µL of RNA gel extraction buffer (0.3 M NaOAc, 1 mM EDTA, 0.25% SDS) was added to each sample, frozen at -80°C for 30 minutes, and then thawed overnight at 22°C while shaking at 700 rpm. Footprints were precipitated by isopropanol and resuspended in 3.5 µL of 10 mM Tris-HCl, pH 8.

### Ribo-seq and Disome-seq library preparation

Library construction was performed as previously described^76,78^: Prior to library construction, barcoded RNA linkers containing a 5 nt unique molecular identifiers (UMIs) were adenylated with Mth RNA ligase (NEB, E2610). Monosome footprints, disome footprints, and size markers were first dephosphorylated with T4 PNK (NEB, M0201) for 1 hour at 37°C followed by linker ligation with truncated T4 RNA ligase 2 (NEB, M0351) to pre-adenylated RNA linkers for 3 h at 22°C. Samples were then treated with 5’ deadenylase (NEB, M0331) and RecJ (Biosearch Technologies, RJ411250) exonuclease to remove unligated linkers. Replicate samples with different linkers were combined and recovered using Oligo Clean and Concentrator (Zymo, D4061) columns and eluted with 10 µL of RNAse-free water. Footprint samples were depleted using siTOOLs riboPOOLs rRNA depletion kit (Galen Lab Supplies, dp-K012-000042). Footprints were recovered using Oligo Clean & Concentrator columns and eluted with 10 µL of RNase-free water. cDNA was generated by first priming with NI-802 (which an additional 2 random nts at the 3’ end of the read) at 65°C for 5 minutes and then by using Superscript III (Invitrogen, 56575) (including 1X First strand buffer, 0.5 mM dNTPs, 5 mM DTT, SUPERase-In) at 55°C for 30 minutes. RNA was degraded by NaOH by incubating at 70°C for 20 minutes and cDNAs were recovered with Oligo Clean & Concentrator columns, eluting with 6 µL of RNase-free water. cDNAs of samples and markers were resolved on a 10% TBE-Urea gel (Bio-Rad, 3450089) and stained with SYBR Gold. cDNAs were excised using size markers as a guide. 500 µL of DNA gel elution buffer (0.3 M NaCl, 1 mM EDTA, 10 mM Tris pH 8) were added to cut gel pieces, frozen for 30 minutes at -80°C and incubated overnight at 22°C, shaking at 700 rpm. Samples were recovered with isopropanol precipitation and resuspended in 15 µL of 10 mM Tris, pH 8. cDNAs were circularized with CircLigase (Epicentre, CL4115K). Pilot and preparative PCRs were carried out using forward and reverse primers listed in Table S1 and Phusion Polymerase (Thermo Fisher, F530S), resolved on 8% native acrylamide gels, and stained with SYBR Gold. The optimal number of PCR cycles that generate a visible band at the correct size without the presence of higher molecular weight bands was determined. Final PCR products were excised from the 8% native acrylamide gel and footprints were eluted with 500 µL of DNA gel elution buffer (0.3 M NaCl, 1 mM EDTA, 10 mM Tris pH 8), and recovered by isopropanol precipitation. Libraries were quantified using TapeStation High Sensitivity D1000 ScreenTape (Agilent, 5067-5584) and sequenced on a NovaSeq X 1.5B chip SE100 by Vanderbilt Technologies for Advanced Genomics (VANTAGE).

### RNA-seq

Total RNA was purified from 100 µL of cell lysate (described above in the “Ribo-seq and Disome-seq lysis, footprinting, and size selection” section) by using TRIzol/chloroform extraction and isopropanol precipitation. GENEWIZ provided Illumina mRNA library preparation and generated 150 bp pair-end reads on Illumina NovaSeq platform.

### Processing of Ribo-seq and Disome-seq data

The data was processed as previously described^79^ using The Advanced Computing Center for Research and Education (ACCRE) at Vanderbilt University: reads were trimmed of adapters and demultiplexed using Cutadapt (v5.1) “cutadapt/5.1 --discard-untrimmed --no-indels -e 0.1 -m 32 -M 41” command for Ribo-seq or “cutadapt/5.1 --discard-untrimmed --no-indels -e 0.1 -m 57 -M 87” for Disome-seq. Reads mapping to rRNAs and tRNAs were removed using Bowtie^80^, with “bowtie/1.3.1 -v 2 -y -S -p 12 --trim5 2 --trim3 5” command. Spike-in reads were aligned to a spike-in FASTA reference using same bowtie parameters. A custom python script was used to remove PCR duplicates based on comparison of 7-nucleotide UMIs. Reads were further trimmed and size selected using “cutadapt/5.1 -u 2 -u -5 -m 25 -M 34” command for Ribo-seq and “cutadapt/5.1 -u 2 -u -5 -m 51 -M 75” for Disome-seq. Reads were then aligned to a revised version of the ribofootPrinter version 1 transcriptome file, using “bowtie/1.3.1 -S -v 1 -y -a -m 1 --best –strata” command.

SAM files were processed using mammalian_builddense.py script, which generates python pickle files containing the sequence and mapped footprint information. After the initial analysis of these files, we found that the first 2 nucleotides of footprints contain mismatches arising from library preparation (not 5-AzaC-induced). To improve the precision of 5-AzaC-induced mismatches, we removed the first 2 nt from the 5’ end of all the reads “bowtie/1.3.1 -S -v 1 -y -a -m 1 --trim5 2 --best --strata”. We also mapped this data to mitochondrial mRNAs using “bowtie -S -v 1 -y -a --trim5 2" command. The cytoplasmic footprint length distribution was obtained with FastQC (v0.12.1) (Babraham Bioinformatics). Footprints were re-processed using the mammalian_builddense.py by choosing 23-32 nt range for monosome and 49-73 nt for disome footprints. Rpkm values were generated for individual genes by dividing the number of reads mapped to a gene by the product of gene length (in kilobases) and the total number of mapped reads (in millions). The gene snapshots were generated using “writegene2” function of Ribofootprinter v1^81^, and plotted with Igor (v9.05, Wavemetrics).

### Processing of RNA-seq data

A STAR index was generated using GRCh38.14 reference genome FASTA and corresponding GTF annotation with --sjdbOverhang set to 99. Paired-end RNA-seq reads were aligned to human genome (hg38) using STAR (v2.7.11b) with “--quantMode GeneCounts” command. Gene level counts were generated using Rsubread (v2.8.1), requiring both read pairs to map, counting only exon reads and grouping by gene ID.

Differential expression analysis was conducted by DESeq2^82^ using raw counts for each gene and running DESeq2 on Rstudio (v2024.04.2+764). Adjusted p-value (padj) <0.05 was used as a significance cutoff. Volcano plots were generated by using ggplot2^83^ in Rstudio. Rpkm values were generated for individual genes by dividing the number of reads mapped to a gene by the product of gene length (in kilobases) and the total number of mapped reads (in millions).

### Mismatch analysis

Ribo-seq and Disome-seq mismatch frequencies were quantified using custom python scripts, “sam_mismatch_analyzer.py” and “sam_mismatch_5prime_distribution.py”. For Ribo-seq data, analyses were performed using “python/3.12.2 sam_mismatch_analyzer.py -d sam_inputs_directory --cds-gtf reference.gtf --offset 12 -o output.xlsx and python/3.12.2 sam_mismatch_5prime_distribution.py -d sam_inputs_directory --cds-gtf reference.gtf --offset 12 -o output.xlsx” command, while Disome-seq data were analyzed using “python/3.12.2 sam_mismatch_analyzer.py -d sam_inputs_directory --cds-gtf reference.gtf --offset 42 -o output.xlsx and python/3.12.2 sam_mismatch_5prime_distribution.py -d sam_inputs_directory --cds-gtf reference.gtf --offset 42 -o output.xlsx” command. In these analyses, reads were classified based on the presence of mismatches, the position of the mismatch within the read, and the identity of the conversion. Only reads with P-sites (offset values of 12 for Ribo-seq and 42 for Disome-seq) located within annotated transcripts were included. Mismatch frequency for each nucleotide was calculated as the number of reads containing a given mismatch divided by the total number of mapped reads. Positional mismatch frequency along the footprint was calculated by dividing the number of reads with a mismatch at a given position by the total number of reads containing that mismatch.

For RNA-seq mismatch analysis, genome-aligned and sorted BAM files were indexed and processed using the custom RNAseq_mismatch_analysis.py script using “RNAseq_mismatch_analysis.py --ref-base C --read-base G --bins 20 --gene-threshold 0.05” command. This script was utilized to quantify the percentage of C-to-G conversions across all reads within each BAM file. For each read, we also determined whether the C-to-G conversion rate is ≥%5. At the gene level, this analysis reported the fraction of reads containing C-to-G conversion rate is ≥%5, providing a measure of gene-specific 5-AzaC incorporation.

### Ribo-seq mismatch vs mRNA half-life analysis

Three biological replicates of mRNA half-life measurements from HCT116 cells were obtained from the Transcriptome Turnover Database^63,84^. The relationship between mismatch frequency and mRNA half-life was analyzed using the custom script “calculate_gene_mismatch_halflife.py” using “python3 calculate_gene_mismatch_halflife.py --sam input.sam --out --offset 12 output.xlsx --gtf input.gtf --half-life-csv half_life_1.csv half_life_2.csv”. Only reads with P-sites (offset value of 12) located within annotated transcripts were included. This algorithm was used to calculate the fraction of the reads containing a mismatch per gene. These values were then plotted against log_10_(half-life) in Prism (v11.0.0). Spearman correlation values were calculated based on the mismatch frequency compared to the log_10_(half-life).

### Ribo-seq and Disome-seq pause analysis

Pause score analysis was performed using “posavgcaller_v2.py” and “tools_m_v2.py” scripts that were modified from posavg function of Ribofootprinter v1^81^. This algorithm generates average pause scores for amino acid or codon motifs by dividing the reads at the motif of interest by the average reads in a designated window (±50 nt of the motif in this study). For computing the pause scores of motifs at A-site, reads were shifted by 14 nt for Ribo-seq and 44 nt for Disome-seq from the 5′ end of the footprint.

Disome peak scores were calculated using “Peak_comparison_v1.py” script with an offset of 42 nt and a cutoff of 2. This algorithm generates the ratio of the peak height at every codon relative to the average codon height for the encompassing gene, excluding sites without peaks. Codons exceeding the specified cutoff in any dataset were reported across all datasets, along with the corresponding nucleotide and amino acid sequences of the A-site, P-site, and the preceding five codons. The resulting disome peak scores were averaged for two biological replicates and the scatter plot was generated in Prism (v11.0.0).

### Ribo-seq and Disome-seq metagene analysis

Metagene analysis was performed using “metagene_transcript.py” adapted from the “metagene” function of Ribofootprinter v1^81^. Reads within the untranslated and coding regions were sorted into 100, 300, and 100 bins, respectively. Read counts in each bin were normalized to total CDS reads per gene, winsorized with a 99^th^ percentile cutoff and averaged across genes. The mean and standard deviation of the two biological replicates from the resulting data were plotted by using Igor (v9.05, Wavemetrics).

---

**Figure S1.**
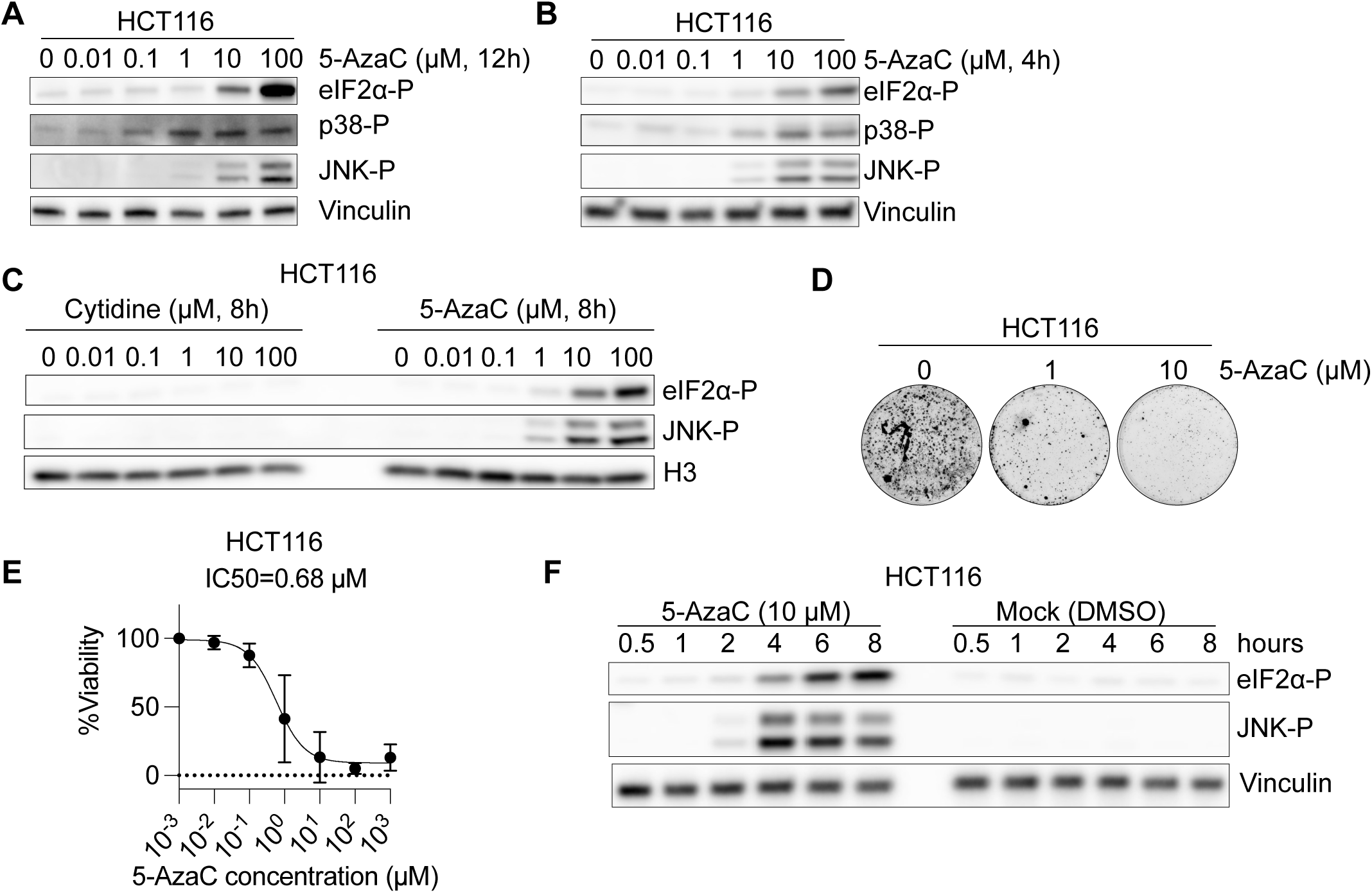
Detailed analysis of 5-AzaC-induced stress signaling. Related to Figure 1. (A) Immunoblots of eIF2α-P, p38-P, JNK-P in HCT116 cells treated with the indicated concentrations of 5-AzaC for 12 h. Vinculin serves as loading control. (B) Immunoblots of eIF2α-P, p38-P, JNK-P in HCT116 cells treated with the indicated concentrations of 5-AzaC for 4 h. Vinculin serves as loading control. (C) Immunoblots of eIF2α-P, and JNK-P in HCT116 cells treated with the indicated concentrations of cytidine or 5-AzaC for 8 h. H3 serves as loading control. (D) Colony formation assay in HCT116 cells with indicated concentrations of 5-AzaC. Biological replicate for the assay in Figure 1F. (E) Viability assay conducted in HCT116 cells that are treated with indicated concentrations of 5-AzaC. IC50 of 5-AzaC is calculated as 0.68 µM. Error bar indicates mean ± standard deviation of three biological replicates. (F) Time course immunoblots of eIF2α-P and JNK-P in HCT116 cells treated with DMSO (Mock) or 10 µM 5-AzaC for 0.5-8 h. Vinculin serves as loading control.

**Figure S2.**
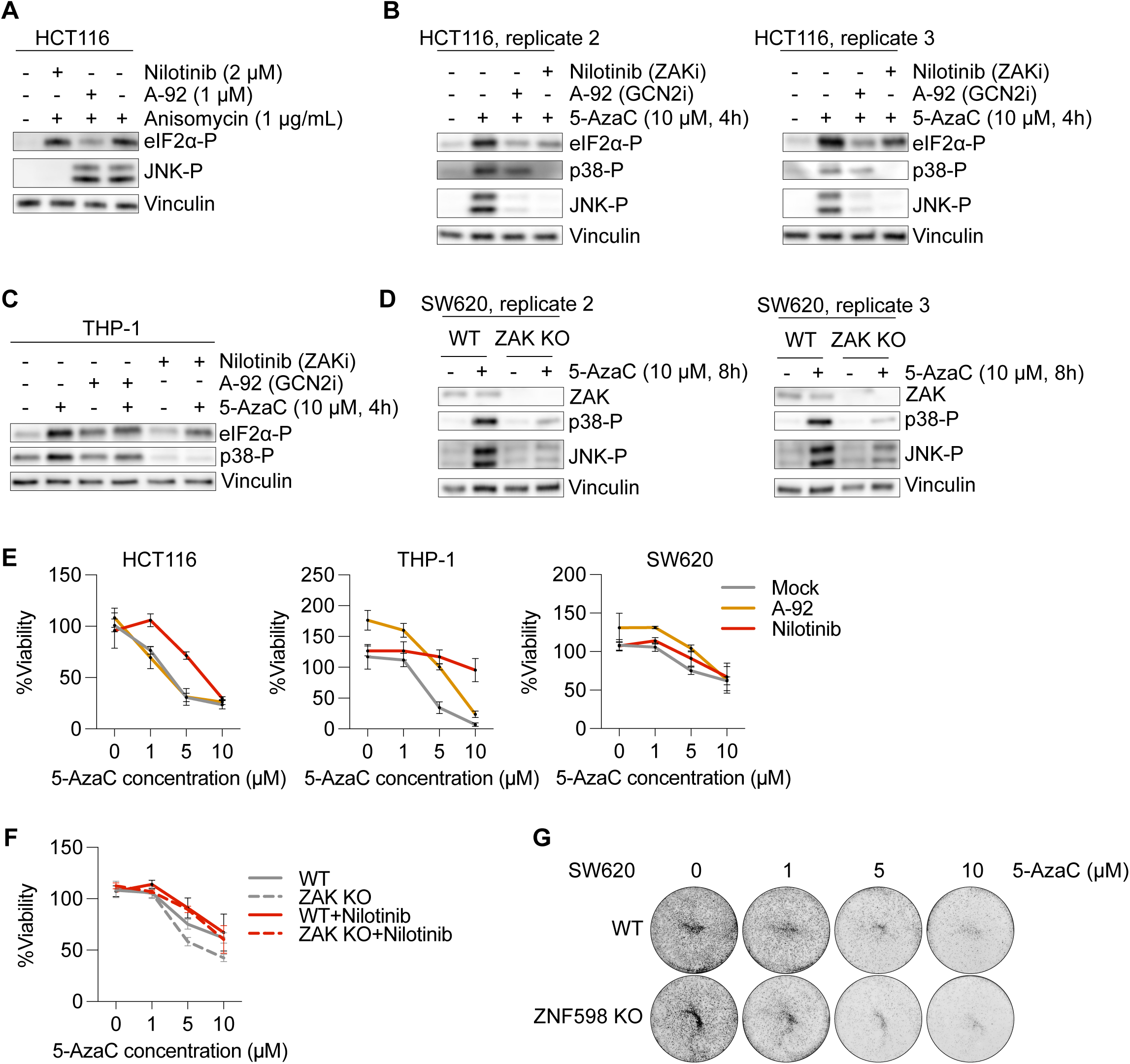
The role of ZNF598, GCN2 and ZAK in 5-AzaC-induced stress signaling. Related to Figure 2. (A) Immunoblots of anisomycin-induced eIF2α-P and JNK-P in HCT116 cells pre-treated with A-92 (2 µM, GCN2 inhibitor) or Nilotinib (1 µM, ZAK inhibitor). 1 µg/mL anisomycin was used to induce disomes. (B) Immunoblots of 5-AzaC-induced eIF2α-P, p38-P, JNK-P in HCT116 cells pre-treated with A-92 (2 µM, GCN2 inhibitor) or Nilotinib (1 µM, ZAK inhibitor). Vinculin serves as loading control. These blots serve as the two other biological replicates of the blot shown in Figure 2A. (C) Immunoblots of 5-AzaC-induced eIF2α-P and p38-P in THP-1 cells pre-treated with A-92 (2 µM, GCN2 inhibitor) or Nilotinib (1 µM, ZAK inhibitor). (D) Immunoblots of ZAK, p38-P and JNK-P in SW620 WT and ZAK KO. Vinculin serves as loading control. These blots serve as the two other biological replicates of the blot shown in Figure 2B. (E) Viability assay measuring the effect of A-92 (orange) and Nilotinib (red) on 5-AzaC-induced cell death in HCT116 (left), THP-1 (middle), and SW620 (right) cells. “Mock” (gray) represents the cells that are treated with 5-AzaC as well as the carrier for A-92 and Nilotinib (DMSO). (F) Viability assay measuring the effect of Nilotinib (red) on 5-AzaC-induced cell death in SW620 WT and SW620 ZAK KO (dashed lines) cells. “Mock” (gray) represents the cells that are treated with 5-AzaC as well as the carrier for Nilotinib (DMSO). (G) Colony formation assay in SW620 WT and ZNF598 KO cells with indicated concentrations of 5-AzaC. Biological replicate of Figure 2D. Where applicable, error bar indicates mean ± standard deviation of 3-9 biological replicates.

**Figure S3.**
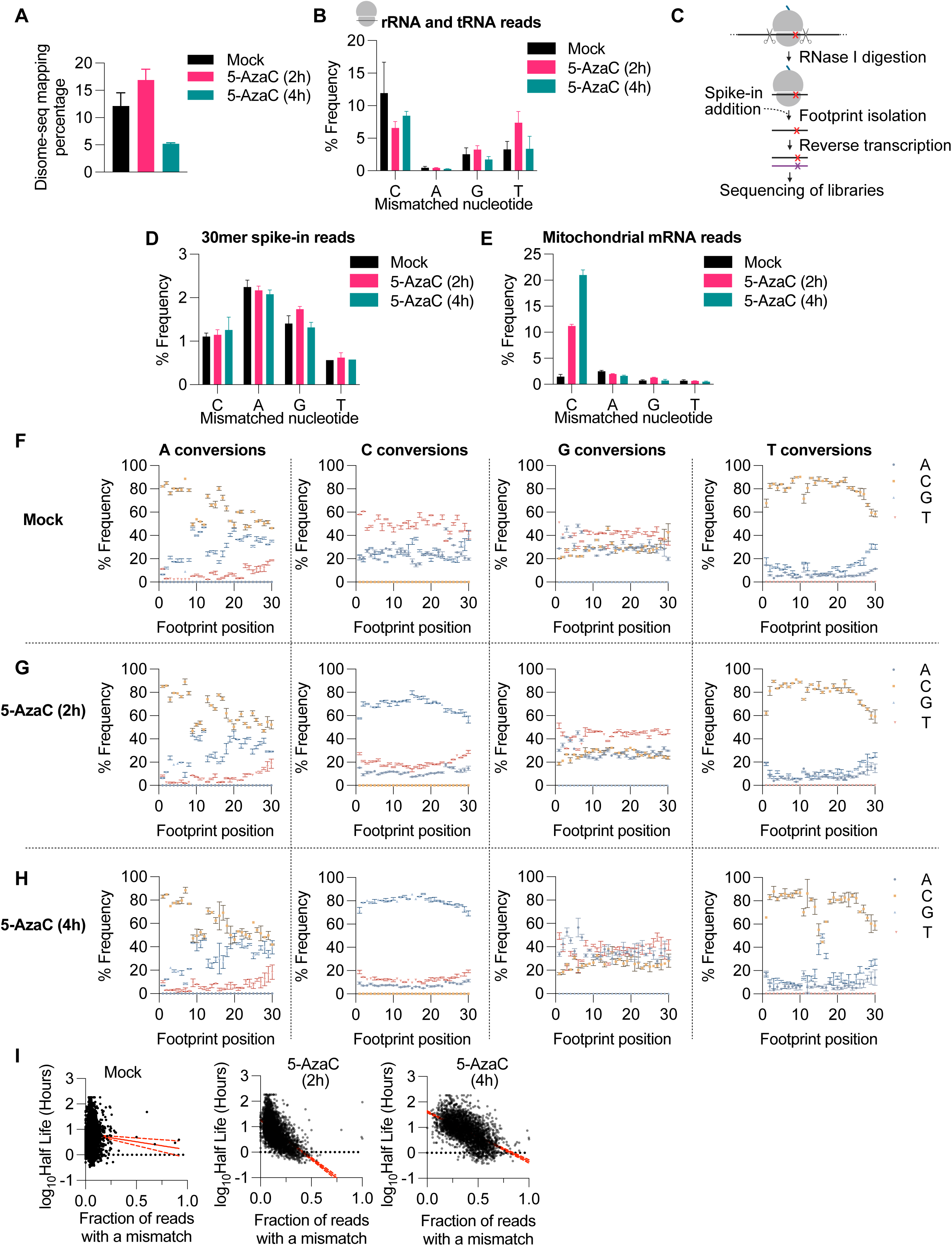
The extent of 5-AzaC-induced C-to-G conversions. Related to Figure 3. (A) Percentage of Disome-seq reads mapping to the transcriptome over the course of 5-AzaC treatment. (B) Nucleotide mismatch frequencies for C, A, G, T in rRNA and tRNA reads in the Ribo-seq data over the course of 5-AzaC treatment. (C) Schematic of our library preparation process. Spike-in addition step, which occurs after footprint isolation and prior to reverse transcription, is indicated with an arrow. (D) Nucleotide mismatch frequencies for C, A, G, T in 30mer spike-in reads. (E) Nucleotide mismatch frequencies for C, A, G, T in mitochondrial mRNA reads. Identity of conversions for A, C, G, and T relative to Ribo-seq footprint position in Mock (F), 5-AzaC (2h) (G) and 5-AzaC (4h) data (H). (I) Replicate data for correlation between mRNA half-life and Ribo-seq mismatch rate in Mock (left), 5-AzaC (2h) (middle), and 5-AzaC (4h) (right). Where applicable, error bar indicates mean ± standard deviation of two biological replicates.

**Figure S4.**
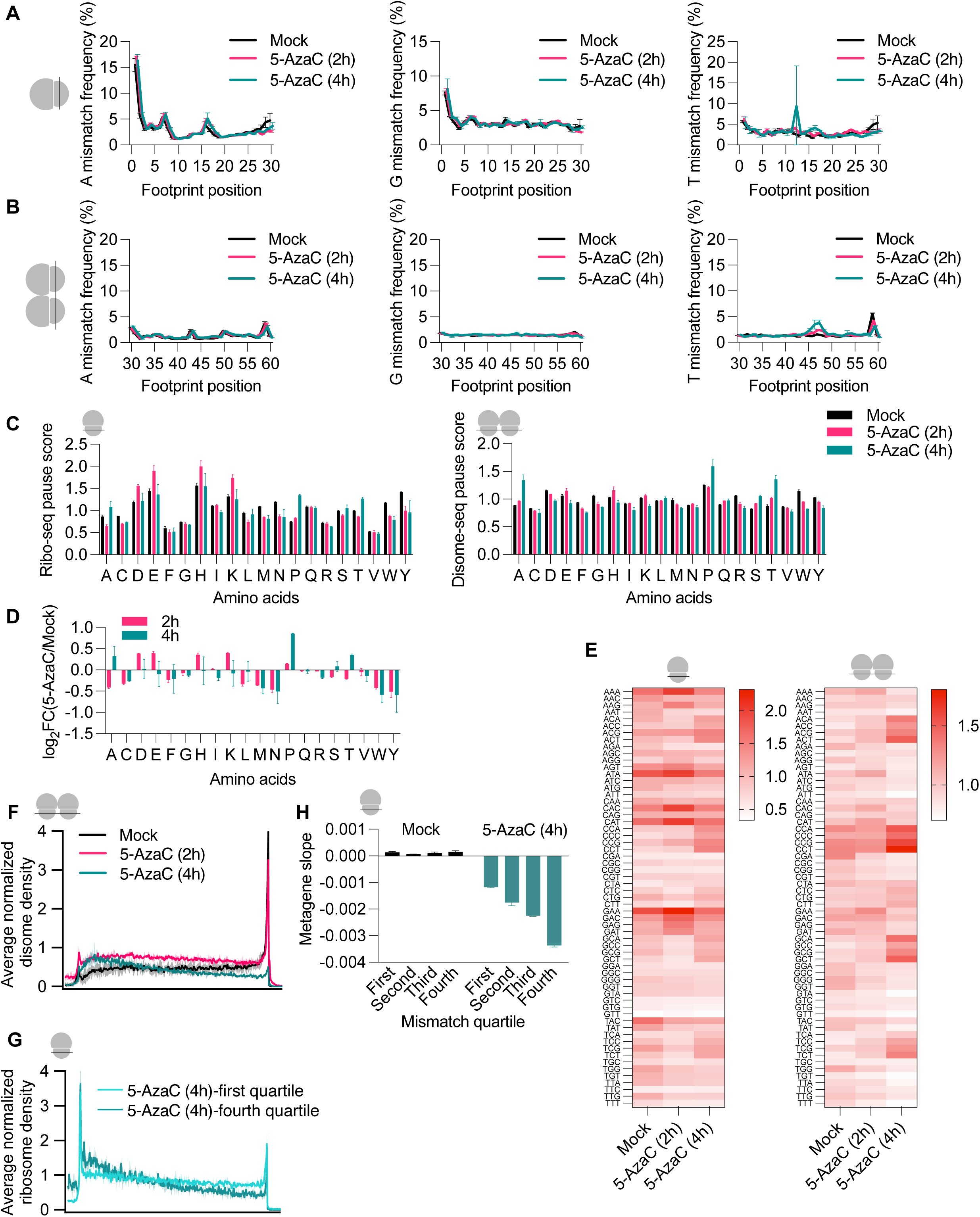
The extent of 5-AzaC-induced C-to-G conversions. Related to Figure 4. (A) A (left), G (middle), and T (right) mismatch frequency across Ribo-seq footprint positions. (B) A (left), G (middle), and T (right) mismatch frequency across Disome-seq footprint positions. We predict that the mild peak with T-mismatch frequency at position 46 may reflect misannotated/mutated T sequences that are actually C and converted to G due to 5-AzaC. (C) Ribo-seq (top) and Disome-seq (bottom) pause scores for all amino acids. (D) Ribo-seq pause score analysis of all amino acids showing 5-AzaC-induced enrichment at Ala, Pro, Ser, Thr codons. FC=Fold Change. Note that the enrichment sequence context is more pronounced for the Disome-seq data in Figure 4D. (E) Ribo-seq (left) and Disome-seq (right) pause score analysis for all the codons showing the codon context for 5-AzaC-induced ribosome stalling and collisions. (F) Disome-seq metagene analysis showing re-distribution of disomes with 4h 5-AzaC treatment, where majority of ribosomes are accumulated at the 5’ of the genes. (G) Ribo-seq metagene profiles of 4h 5-AzaC treatment. The data was first binned to four quartiles based on mismatch rate per gene (first quartile being the lowest and fourth quartile being the highest mismatch), and then metagene analysis of each quartile was conducted. This figure compares the first and fourth quartiles, latter of which shows the unique 3’ depletion of reads. (H) Ribo-seq metagene slope measurement of each quartile for Mock vs 4h 5-AzaC treatment. Slope of the values corresponding to coding regions was calculated for each quartile and was plotted. 5-AzaC treatment shows mismatch rate-dependent increase of the negative slope. Where applicable, error bar indicates mean ± standard deviation of two biological replicates.

