## Supplementary material for "5-Azacytidine incorporation into mRNAs disrupts translation and induces ribosome collisions": Table S2

Table S2. Ribo-seq, Disome-seq, and RNA-seq statistics.

| ID | Description | Data type | Reads aligned to rRNA/tRNA | Not aligned to rRNA/tRNA | Reads without PCR duplicates | Aligned to transcriptome | Aligned to genome | Aligned to mitochondria transcriptome |
|---|---|---|---|---|---|---|---|---|
| JM011F | Mock-1 | Ribo-seq | 5807217 | 20578113 | 8075788 | 4485840 | | 25604 |
| JM012F | Mock-2 | Ribo-seq | 10179867 | 34060143 | 13787616 | 7884823 | | 39818 |
| JM013F | 5-AzaC (2h)-1 | Ribo-seq | 5090265 | 19716025 | 6989957 | 3210459 | | 82913 |
| JM014F | 5-AzaC (2h)-2 | Ribo-seq | 6872403 | 21775897 | 8765636 | 4015501 | | 129714 |
| JM015F | 5-AzaC (4h)-1 | Ribo-seq | 7179285 | 25618884 | 5730247 | 1503755 | | 32331 |
| JM016F | 5-AzaC (4h)-2 | Ribo-seq | 5878460 | 24383442 | 5290747 | 1064536 | | 25397 |
| JM011Fd | Mock-1 | Disome-seq | 50643692 | 28096912 | 5554208 | 578109 | | |
| JM012Fd | Mock-2 | Disome-seq | 118998682 | 51833421 | 9842132 | 1360442 | | |
| JM013Fd | 5-AzaC (2h)-1 | Disome-seq | 18878521 | 45705844 | 11951515 | 1847506 | | |
| JM014Fd | 5-AzaC (2h)-2 | Disome-seq | 22894582 | 56138270 | 17493703 | 3202467 | | |
| JM015Fd | 5-AzaC (4h)-1 | Disome-seq | 67536294 | 36849574 | 9650540 | 487412 | | |
| JM016Fd | 5-AzaC (4h)-2 | Disome-seq | 26551090 | 30397667 | 8089073 | 430844 | | |
| JM011M | Mock-1 | RNA-seq | | | | | 42020091 | |
| JM012M | Mock-2 | RNA-seq | | | | | 38582613 | |
| JM013M | 5-AzaC (2h)-1 | RNA-seq | | | | | 43425939 | |
| JM014M | 5-AzaC (2h)-2 | RNA-seq | | | | | 37468140 | |
| JM015M | 5-AzaC (4h)-1 | RNA-seq | | | | | 41437696 | |
| JM016M | 5-AzaC (4h)-2 | RNA-seq | | | | | 33749167 | |
