## Supplementary material for "5-Azacytidine incorporation into mRNAs disrupts translation and induces ribosome collisions": Table S1

Table S1. Oligonucleotides used in this study.

| Name | Sequence |
|---|---|
| **Oligos used for cloning lentiGuide-Puro plasmids** | |
| ZAK-pLentiguide-Fwd | 5’-CACCGTGTATGGTTATGGAACCGAG-3’ |
| ZAK-pLentiguide-Rev | 5’-AAACCTCGGTTCCATAACCATACAC-3’ |
| ZNF598-pLentiguide-Fwd | 5’-CACCGACCGCTGCTCTACCAAGATG-3’ |
| ZNF598-pLentiguide-Rev | 5’-AAACCATCTTGGTAGAGCAGCGGTC-3’ |
| **Ribo-seq/Disome-seq size selection markers** | |
| 25mer | rArUrGrUrArCrArCrGrGrArGrUrCrGrArGrCrArCrCrCrGrCrA |
| 34mer | rArUrGrUrArCrArCrGrGrArGrUrCrGrArGrCrArCrCrCrGrCrArArCrGrCrGrArArUrG |
| 80mer | rArUrGrUrArCrArCrGrGrArGrUrCrGrArGrCrArCrCrCrGrCrArArCrGrCrGrArArUrGrUrAr CrArCrGrGrArGrUrCrGrArGrCrArCrCrCrGrCrArArCrGrCrGrArUrGrUrArCrArCrCrCrGr CrArArCrGrCrGrA |
| **Spike-in oligos** | |
| 30mer-1 | rArArUrArCrCrArCrCrCrCrCrArUrGrArArCrGrCrUrGrCrArCrArCrArCrG |
| 30mer-2 | rArArCrUrArCrCrGrArCrUrCrArUrCrCrCrArUrCrUrUrGrCrCrArGrUrArC |
| 30mer-3 | rCrUrArArUrArCrUrUrArCrGrArArCrCrArGrArCrGrArArUrCrCrCrUrUrG |
| 60mer-1 | rArArUrArCrCrArCrCrCrCrCrArUrGrArArCrGrCrUrGrCrArCrArCrArCrGrArArUrArCrCr ArCrCrCrCrCrArUrGrArArCrGrCrUrGrCrArCrArCrArCrG |
| 60mer-2 | rArArCrUrArCrCrGrArCrUrCrArUrCrCrCrArUrCrUrUrGrCrCrArGrUrArCrArArCrUrArCr CrGrArCrUrCrArUrCrCrCrArUrCrUrUrGrCrCrArGrUrArC |
| 60mer-3 | rCrUrArArUrArCrUrUrArCrGrArArCrCrArGrArCrGrArArUrCrCrCrUrUrGrCrUrArArUrAr CrUrUrArCrGrArArCrCrArGrArCrGrArArUrCrCrCrUrUrG |
| **Ribo-seq/Disome-seq Linkers** | |
| NI-810 | 5´-/5Phos/NNNNNATCGTAGATCGGAAGAGCACACGTCTGAA/3ddC/ |
| NI-811 | 5´-/5Phos/NNNNNAGCTAAGATCGGAAGAGCACACGTCTGAA/3ddC/ |
| NI-812 | 5´-/5Phos/NNNNNCGTAAAGATCGGAAGAGCACACGTCTGAA/3ddC/ |
| NI-813 | 5´-/5Phos/NNNNNCTAGAAGATCGGAAGAGCACACGTCTGAA/3ddC/ |
| NI-814 | 5´-/5Phos/NNNNNGATCAAGATCGGAAGAGCACACGTCTGAA/3ddC/ |
| NI-815 | 5´-/5Phos/NNNNNGCATAAGATCGGAAGAGCACACGTCTGAA/3ddC/ |
| **Ribo-seq/Disome-seq RT primer** | |
| NI-802 | 5'-/5Phos/NNAGATCGGAAGAGCGTCGTGTAGGGAAAGAG/iSp18/GTGACTGGAGTT CAGACGTGTGCTC |
| **Ribo-seq/Disome-seq Library PCR primers** | |
| NI-NI-798 | 5´-AATGATACGGCGACCACCGAGATCTACACTCTTTCCCTACACGACGCTC |
| NI-822 | 5’- CAAGCAGAAGACGGCATACGAGATACATCGGTGACTGGAGTTCAGACGTGTG |
| NI-823 | 5’-CAAGCAGAAGACGGCATACGAGATGCCTAAGTGACTGGAGTTCAGACGTGTG |
